# Polarly Localized WPR Proteins Interact With PAN Receptors And The Actin Cytoskeleton During Maize Stomatal Development

**DOI:** 10.1101/2022.04.22.489215

**Authors:** Qiong Nan, Si Nian Char, Bing Yang, Eric J. Bennett, Bing Yang, Michelle R. Facette

**Author notes:** The author responsible for distribution of materials integral to the findings presented in this article in accordance with the policy described in the Instructions for Authors is: Michelle Facette.

## Abstract

Polarization of cells prior to asymmetric cell division is crucial for correct cell divisions, cell fate and tissue patterning. In maize stomatal development, polarization of subsidiary mother cells prior to asymmetric division is controlled by the BRK-PAN-ROP pathway. Two catalytically inactive receptor-like kinases, PAN2 and PAN1, are required for correct division plane positioning. Proteins in the BRK-PAN-ROP pathway are polarized in subsidiary mother cells, with the polarization of each protein dependent on the previous one. As most of the known proteins in this pathway do not physically interact, possible interactors that might participate in the pathway are yet to be described. We identified WPR proteins as new players during subsidiary mother cell polarization. WPRs physically interact with PAN receptors, and polarly accumulate in subsidiary mother cells. The polarized localization of WPR proteins depends on PAN2 but not PAN1. CRISPR-Cas9- induced mutations result in division plane defects in subsidiary mother cells, and ectopic expression of WPR-RFP results in stomatal defects and alterations to the actin cytoskeleton. We show certain WPR proteins directly interact with F-actin through their N-terminus. Our data implicate WPR proteins as potentially regulating actin filaments, which providing insight into their molecular function. Together, these results demonstrate that WPR proteins are important for cell polarization.

**One-sentence summary:** Four related proteins, identified via their physical interaction with the receptor PAN2, are polarly localized prior to asymmetric division in stomatal lineage cells, and interact with F-actin.

## INTRODUCTION

Asymmetric cell division, which generates two daughter cells with different fates, is indispensable for cellular differentiation and diversity. In plants, cells do not move, therefore asymmetric cell division is particularly important for cellular patterning (De Smet and Beeckman, 2011; Muroyama and Bergmann, 2019). Certain processes are conserved during asymmetric cell division, although the specific molecular players may vary. A key principle of asymmetric cell division is cell polarization prior to mitosis, which is often marked by the polarization of plasma membrane–associated proteins and nuclear migration to the future division plane (Facette and Smith, 2012; Galatis and Apostolakos, 2004; Yoshida et al., 2019). Cell polarization may be determined by intrinsic cues (pre-existing within the mother cell) or extrinsic cues (originating from outside the mother cell) (Yang 2008; Facette et al., 2019; Lipka et al., 2015).

The asymmetric division of maize subsidiary mother cells (SMCs) has been used as a model to understand cell polarization and asymmetric division in plants. Grasses such as *Zea mays* (maize), *Oryza spp* (rice) and *Brachypodium distachyon* (purple false brome) possess 4-celled stomatal complexes consisting of two guard cells flanked by a pair of subsidiary cells **(Supplemental Figure 1)**. Grass stomatal formation is initiated by an asymmetric division that generates a guard mother cell (GMC). The lateral neighbors of the GMC, called subsidiary mother cells (SMCs), divide asymmetrically to produce subsidiary cells that flank the GMC. It is proposed that SMC divisions are induced by an extrinsic cue from the GMC. Eventually, the GMC divides symmetrically to form a pair of guard cells flanked by the subsidiary cells (Stebbins and Shah, 1960; Facette and Smith, 2012; Raissig et al., 2017; Nunes et al., 2020; Gray et al., 2020).

In maize SMCs, several plasma membrane–associated proteins make up the BRK-PAN-ROP pathway, which promotes SMC polarity. BRICK1 (BRK1) is the earliest marker of SMC polarity (Facette et al., 2015). BRICK1 is a member of SCAR/WAVE complex that promotes branched actin networks via activation of the Arp2/3 complex (Frank and Smith 2002; Facette et al., 2015). BRK1, and all subsequent proteins, are polarized within SMCs, accumulating at the site of GMC contact. BRK1 is required for PANGLOSS2 (PAN2) and PAN1 polarization, which are two leucine-rich repeat receptor-like proteins with inactive kinase domains. PAN2 is required for the subsequent polarization of PAN1 (Zhang et al., 2012). Plant Rho family GTPase (ROP) is a small GTPase that physically interacts with PAN1 and its polarization is dependent on PAN1 (Humphries et al., 2011). After the polarized accumulation of these proteins, an actin patch appears at the SMC-GMC interface and the pre-mitotic SMC nucleus migrates towards the GMC. Within the pathway, only PAN1 and ROP have been shown to physically interact, however each protein is required for the next to be polarized. This implies that there are additional proteins important for the recruitment and/or polarization of each of the known players. Moreover, the molecular role that each of these proteins performs during SMC polarization is not known, including how polarization of these membrane-localized proteins culminates in nuclear migration and positioning of the division plane.

Since PAN1 and PAN2 are catalytically inactive kinases (Humphries et al., 2011; Zhang et al., 2012), it is unclear what their molecular function is. They may act as pseudokinases with important roles in scaffolding; however other than ROP no other PAN-interacting proteins have yet been characterized. In some cases, catalytically inactive receptors partner with active leucine-rich repeat receptor kinases and act as inhibitors of function (Lee et al., 2012), but to date no active kinases have been identified in the pathway. PAN1 and PAN2 might perform functions similar to other polarized plasma membrane proteins and mediate diverse downstream functions that relate to cellular polarization such as: signal transduction, accumulation of other cell polarity factors, or eventual nuclear migration (De Smet and Beeckman, 2011; Ashraf and Facette, 2020). For example, in *A. thaliana* stomatal development, the polarized proteins BREAKING OF ASYMMETRY IN THE STOMATAL LINEAGE (BASL) and POLAR LOCALIZATION DURING ASYMMETRIC DIVISION AND REDISTRIBUTION (POLAR) act as scaffolding proteins for localized signal transduction (Dong et al., 2009; Pillitteri et al., 2011; Houbaert et al., 2018). Notably, the *B. distachyon* orthologue of POLAR was recently discovered to have a distinct localization pattern during grass stomatal development that is opposite that of PAN1 – it is excluded from the GMC-SMC contact site (Zhang et al., 2022). Polarized BASL is required for pre-mitotic microtubule-based nuclear migration and post-mitotic actin-based nuclear migration (Muroyama et al., 2020). The LRR-RLK INFLORESCENCE AND ROOT APICES RECEPTOR KINASE (IRK) is polarly localized in root cells that divide (or have recently divided) asymmetrically however the molecular pathway downstream of IRK is not characterized (Campos et al., 2020).

Polarized accumulation of actin filaments is a prominent feature of maize SMC division, but the functional role of the actin patch that forms late in SMC polarization is unclear. While SMC nuclear migration in grasses is based on actin networks (Cho and Wick, 1990), formation of the actin patch and nuclear migration can be uncoupled (Cartwright et al., 2009; Apostolakos et al., 2018), implying that the actin patch has a function distinct from promoting nuclear migration. It is plausible that ROP GTPases are important for stimulating actin patch formation, however this has not been clearly demonstrated. In addition to the actin patch, there are other important actin-related processes during SMC polarization. The early role of BRK proteins, actin-mediated nuclear migration and actin patch formation imply multiple functions for actin networks in SMC polarization. SMC actin filaments may provide force or/and tracks for polarized organelle movement, and/or may facilitate endo/exocytosis at the polarity site (Hadley et al., 2006; Kimata et al., 2016; Wu and Bezanilla, 2018).

Actin-dependent movement of chloroplasts has been shown to involve WEAK CHLOROPLAST MOVEMENT UNDER BLUE LIGHT 1 (WEB1) and PLASTID MOVEMENT IMPAIRED 2 (PMI2) (Luesse et al., 2006; Kodama et al., 2010). WEB1, PMI2 and similar WEB1/PMI2-RELATED (WPR) proteins have a DUF827 domain which is a series of coiled-coils; *A. thaliana* has fourteen DUF827-containing proteins including the WEB1-like (WEL) clade, PMI2-LIKE (PMI) clade and the WPRA and WPRB clades (Kodama et al., 2011). Previous studies in *A. thaliana* suggest proteins with this domain have diverse functions. WEB1 (Weak Chloroplast Movement under Blue Light 1) and PMI2 (Plastid Movement Impaired 2) are interacting proteins and are required for actin-based chloroplast photorelocation (Luesse et al., 2006; Kodama et al., 2010). TOUCH-REGULATED PHOSPHOPROTEIN1 (TREPH1) belongs to the WPRA protein clade, and is phosphorylated in response to touch (Wang et al., 2018). TREPH1 is required for touch-induced growth repression. WEB1 is membrane-associated and PMI2 is cytosolic in protoplasts, but *in planta* localization for DUF827 proteins has not been demonstrated. Moreover, while the demonstrated phenotypes of WEB1 and PMI2 implies an actin-related function, no evidence for physical or regulatory interactions with actin have been shown for any DUF827 protein thus far.

In this study, we identified members of the DUF827 domain-containing WPR protein family that participate in maize SMC polarization. WPRs interact with both PAN2 and PAN1, and WPRs can also form homodimers and heterodimers. WPRs localize polarly in SMC at sites of guard mother cell (GMC) contact, and their polarized localization depends on PAN2 but not PAN1. These findings suggest WPRs are plasma membrane-associated proteins that are important for pre-mitotic polarity in maize SMCs, and are the first physical link between PAN1 and PAN2. CRISPR-Cas9- induced *wprb1;wprb2* double mutants and plants ectopically expressing WPRB2 have an increased frequency of aberrantly formed subsidiary cells. WPRB2 overexpression causes a decrease in cellular F-actin. Additionally, we show that WPRB2 directly interacts with F-actin through its N-terminus. In total, these results suggest the WPRs are components of the BRK-PAN-ROP pathway and likely act downstream of PAN2 to regulate actin-associated processes during cell polarization.

## RESULTS

### WPRs Exhibit Polarized Localization in Maize SMCs

To better understand the functions of PAN and BRK proteins during SMC polarization, we identified proteins that physically interact with PAN2-YFP and PAN1-YFP. Co-immunoprecipitation/mass spectrometry (Co-IP/MS) of BRK1-CFP, Rab11D-YFP, PIN1-YFP, PDI-YFP and non-transgenic controls was previously performed using anti-GFP beads (Facette et al., 2015). In parallel, co-IP/MS of PAN2-YFP and PAN1-YFP was also performed, but not presented in Facette et al. (2015). As previously published, a WD-score was calculated to identify high-confidence interactors (Facette et al., 2015; Sowa et al., 2009). Amongst the PAN2-YFP interactors were four related proteins containing Domain of Unknown Function 827 (DUF827) **(Supplemental Dataset 1).** The DUF827 family includes previously identified WEB1-PMI2 RELATED (WPR) proteins (Kodama et al., 2010; Gardiner et al., 2011; Kodama et al., 2011). We identified sixteen WPRs in *A. thaliana* and seventeen WPRs in maize. A protein tree was inferred, and these **(Supplemental Figure 2)** proteins fall into five clades: WPRA, PMI, WEB/WEL, WPRB and WPRC. The four maize proteins interacting with PAN2 fall into WPRA and WPRB clades; we named these proteins WPRA1, WPRA2, WPRB1, and WPRB2. WPRs have predicted coiled-coil domains in their central plant-specific DUF827 region, flanked by uncharacterized N- and C-terminal regions.

Thus far, all proteins identified in the BRK-PAN-ROP pathway polarly localize in SMCs prior to division (Cartwright et al., 2009; Humphries et al., 2011; Zhang et al., 2012; Facette et al., 2015). To determine if WPR proteins play a role in SMC polarization, we determined if they were similarly polarized. We created stable transgenic maize lines expressing either CFP-WPRA2 or RFP-WPRB2. Genomic fragments containing the native promoter, introns and terminator were used. At early stages of stomatal development prior to SMC formation, CFP-WPRA2 **(Figure 1)** and RFP-WPRB2 **(Figure 1B)** predominantly localize to the cell periphery in all cell types, with low levels of cytoplasmic fluorescence. RFP-WPRB2 is enriched in cell corners, similar to BRK1-CFP (Facette et al., 2015). As development proceeds, CFP-WPRA2 and RFP-WPRB2 become enriched in the SMC at the site of GMC contact and remain polarized after SMC division **(Figure 1)**. Plasmolysis experiments confirm that CFP-WPRA2 and RFP-WPRB2 are polarized in the SMC, not the GMC **(Supplemental Figure 3)**. We further validated the localization by performing immunofluorescence using custom-generated antibodies that recognize either WRPA1/WPRA2 or WPRB2 (**Supplemental Fig 4**). Similar to the transgenic lines, we saw polarized localization in SMCs and enrichment at cell corners. Thus, the localization of WPRA2 and WPRB2 is similar to known BRK-PAN-ROP pathway components and supports a role in SMC polarization. Notably, WPR proteins contain no predicted transmembrane domains or lipidation signals, indicating their association with the membrane is either peripheral or via interactions with other membrane proteins.

**Figure 1.**
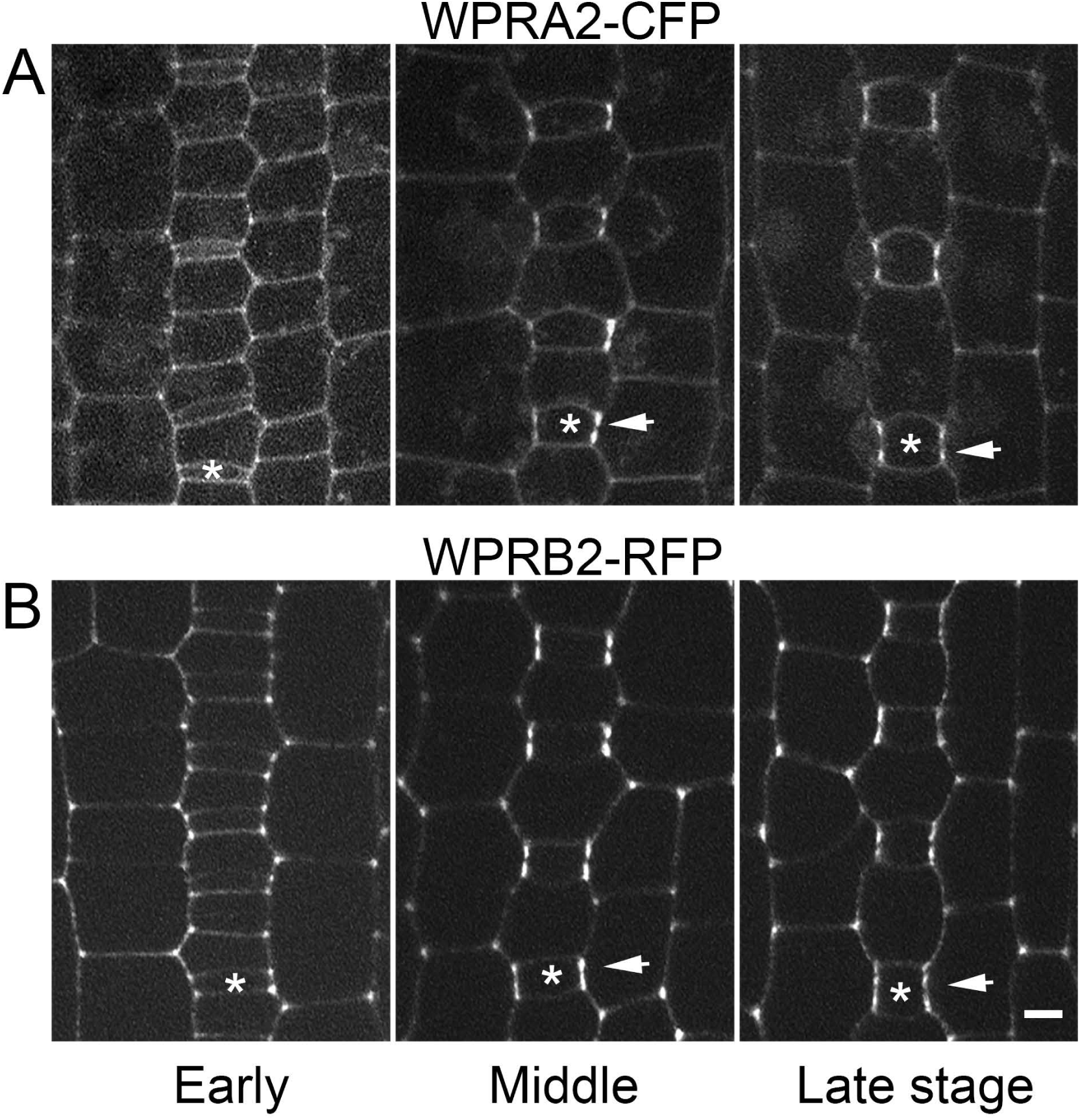
CFP-WPRA2 and RFP-WPRB2 polarize in developing subsidiary mother cells. The stomatal division zone of the leaf epidermis of transgenic maize plants expressing fluorescent fusion proteins was dissected and analyzed. Three different developmental stages were selected according to GMC width and SMC division status to observe the localization of CFP-WPRA2 (A) and RFP-WPRB2 (B). Arrows point to sites of CFP-WPRA2 or RFP-WPRB2 accumulation in SMCs where they contact the adjacent GMC. Asterisks mark a GMC within each stomatal row. Scale bar = 5 µm

### WPRs Interact with PAN1 and PAN2

To confirm the WPR-PAN2 interaction, we performed reciprocal co-IP/MS experiments using three strategies, using either our fluorescent protein fusion lines or a native antibody. Firstly, we used anti-GFP to immunoprecipitate WPRA2 from transgenic plants expressing CFP-WPRA2; non-transgenic wild type plants were used as a negative control. Secondly, we used anti-RFP to immunoprecipitate WPRB2 from transgenic plants expressing RFP-WPRB2, again using non-transgenic plants as a negative control. Thirdly, we used an antibody raised against WPRA1/A2 (see Methods, and **Supplemental Fig. 4**) to pull down endogenous WPRA from non-transgenic plant extracts, with reactions omitting this antibody as a negative control. In each experiment, three biological replicates were performed. After comparing relative abundances of identified proteins with corresponding negative controls, we found WPRA family proteins co-precipitated with WPRB family proteins in all three assays (**Table 1 and Supplement Data Set 2**). Moreover, we found that both PAN1 and PAN2 co-precipitated with WPRA1/A2 and PAN2 co-precipitated with RFP-WPRB2. In addition, enrichment of WPRB3 peptides was detected in CFP-WPRA2 and WPRA1/A2 immunoprecipitates, suggesting WPRB3 may also associate with WPRA proteins.

**Table 1.** WPR proteins interact with other WPR family members and PAN receptors. WPR-interacting proteins identified by Co-IP/MS. For a complete list of all proteins, see Supplementary Data Set 1. Three different experiments were performed. 1: Anti-GFP was used to pull down WPRA2 and interacting proteins from extracts of plants expressing CFP-WPRA2, and from non-transgenic plants as a negative control. 2: Anti-WPRA1/A2 was used to pull down WPRA1/A2 and interacting proteins from B73 extracts; negative controls used the same extracts but omitted the anti-WPRA1/A2 antibody. 3: Anti-RFP was used to pull down WPRB2 and interactors from extracts of plants expressing RFP-WPRB2, and from non-transgenic plants as a negative control. B73 vs4 gene codes are used to identify proteins, and the number of unique peptides (# peptides) is listed. The baits WPRA2, WPRA1/A2, and WPRB2 are highlighted in bold.

| Gene ID |  | Gene name | Rep1<br># Unique<br>Peptides | Rep2<br># Unique<br>Peptides | Rep3<br># Unique<br>Peptides | Control 1<br># Unique<br>Peptides | Control 2<br># Unique<br>Peptides | Control 3<br># Unique<br>Peptides |
| --- | --- | --- | --- | --- | --- | --- | --- | --- |
| Anti-GFP | Zm00001d041088 | **WPRA2 (bait)** | 28 | 42 | 2 | 2 | 0 | 0 |
|  | Zm00001d007164 | WPRB2 | 5 | 3 | 0 | 0 | 0 | 0 |
|  | Zm00001d010610 | WPRB3 | 15 | 13 | 0 | 0 | 0 | 1 |
|  | Zm00001d029420 | WEB subfamily protein | 5 | 7 | 0 | 0 | 0 | 0 |
| Anti-WPR A1/A2 | Zm00001d023629 | **WPRA1 (bait)** | 24 | 19 | 26 | 0 | 1 | 2 |
|  | Zm00001d041088 | **WPRA2 (bait)** | 26 | 19 | 27 | 0 | 0 | 0 |
|  | Zm00001d047516 | WPRB1 | 5 | 4 | 6 | 0 | 0 | 0 |
|  | Zm00001d007164 | WPRB2 | 19 | 21 | 21 | 0 | 0 | 0 |
|  | Zm00001d010610 | WPRB3 | 24 | 21 | 24 | 0 | 1 | 0 |
|  | Zm00001d026139 | WPRC subfamily protein | 2 | 1 | 0 | 0 | 0 | 0 |
|  | Zm00001d000443 | WPRC subfamily protein | 2 | 2 | 0 | 0 | 0 | 1 |
|  | Zm00001d048523 | WEB subfamily protein | 2 | 2 | 2 | 0 | 0 | 0 |
|  | Zm00001d007862 | PAN2 | 4 | 5 | 4 | 0 | 0 | 2 |
|  | Zm00001d031437 | PAN1 | 2 | 5 | 2 | 0 | 0 | 0 |
| Anti-RFP | Zm00001d007164 | **WPRB2 (bait)** | 18 | 20 | 17 | 0 | 1 | 0 |
|  | Zm00001d023629 | WPRA1 | 18 | 16 | 16 | 0 | 0 | 0 |
|  | Zm00001d041088 | WPRA2 | 15 | 21 | 20 | 0 | 2 | 0 |
|  | Zm00001d010610 | WPRB3 | 1 | 2 | 1 | 0 | 0 | 0 |
|  | Zm00001d007826 | PAN2 | 3 | 0 | 1 | 0 | 0 | 0 |

To validate these interactions, and test whether WPRs directly interact with PAN1 and PAN2, yeast two-hybrid assays were conducted using intracellular portions of PAN1 and PAN2 as bait (Zhang et al., 2012), and WPRA1, WPRA2, WPRB1, WPRB2, and WPRB3 as prey. In these assays, no interaction between WPRA1 or WPRA2 and PAN1 or PAN2 was observed. However, we found that both WPRB1 and WPRB2 interact with PAN1 and PAN2. WPRB3 interacts with PAN2 but not PAN1 **(Figure 2A and 2B)**. These data confirm a direct physical interaction between the intracellular regions of PAN and WPRB proteins. Our co-IP MS data suggest WRPA-WPRB interact, and previous work has shown that *A. thaliana* WEB1 interacts with and PMI2 and itself (Kodama et al., 2010). We performed yeast 2-hybrid assays to determine if maize WPR proteins also form homo/heterodimers, and found several hetero- and homo-dimer combinations **(Figure 2C and 2D)**. This includes heterodimers between WPRA and WPRB proteins. This potentially explains why the Co-IP/MS data indicate that PAN proteins interact with both WPRA and WPRB proteins, yet no direct interaction between WPRA and PAN proteins was observed. Taken together, the Co-IP/MS combined with Y2H assays demonstrated an interaction network **(Figure 2F).** We suggest that WPRB could be mediating a WPRA-WPRB heterodimer interaction with both PAN1 and PAN2 **(Figure 2G)**.

**Figure 2.**
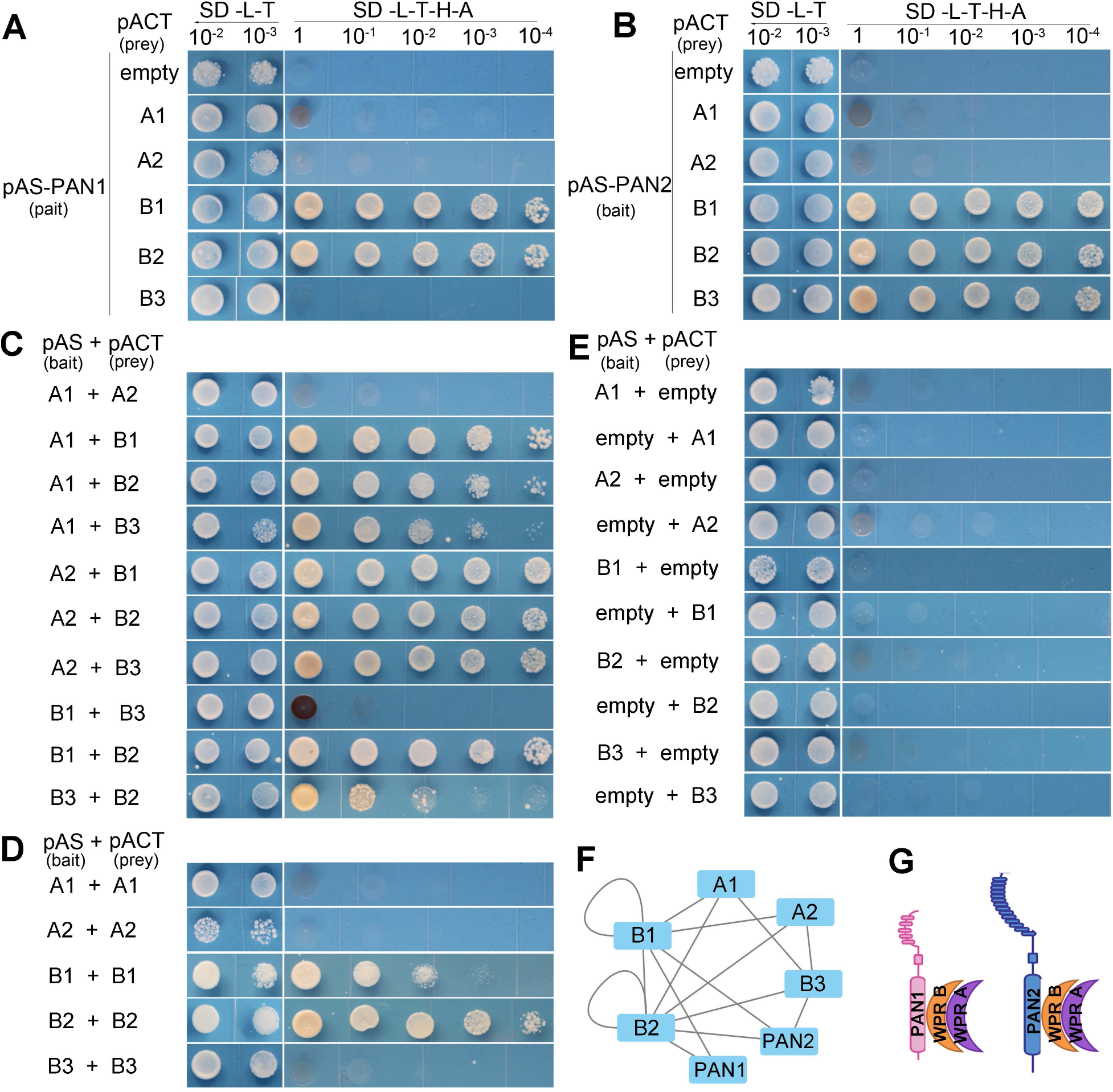
Yeast 2-hybrid analysis of WPR and PAN protein interactions. Interactions were assessed using the GAL4-based yeast two-hybrid system. Yeast was grown at different dilutions on non-selective (-L-T) and selective (-L-T-A-H) media. Soluble intracellular regions of PAN1 and PAN2 were used. **(A)** and **(B)**, interactions of PAN1 and PAN2 with WPRA1, WPRA2, WPRB1, WPRB2, and WPRB3. **(C)** Heterodimer formation was assessed between WPRA1, WPRA2, WPRB1, WPRB2, and WPRB3. **(D)** Homodimer formation was assessed between WPRA1, WPRA2, WPRB1, WPRB2, and WPRB3. **(E)** Negative controls using empty bait plasmid pASGW-attR or prey plasmid pACTGW-attR. **(F)** Network diagram showing the observed interactions between WPRs and PAN1/PAN2. **(G)** A model of protein complex formation between PAN and WPR proteins.

### WPR Proteins Act after PAN2 but before PAN1 in the BRK-PAN-ROP Pathway

Grass stomatal development occurs sequentially, where cells closest to the leaf base are developmentally less advanced, and cells more distal from the leaf base are more developmentally advanced. Therefore, in a single leaf, all stages of development can be observed **(Supplemental Figure 1A-F).** SMCs are initially unpolarized. As development proceeds, proteins important for cell polarization become enriched at the GMC-SMC contact site, in sequence. Previous studies have demonstrated that BRK-PAN-ROP pathway components polarize sequentially in SMCs the following order: BRK1, PAN2, PAN1, followed by formation of a cortical F-actin patch at the GMC contact site (Cartwright et al., 2009; Zhang et al, 2012; Facette et al., 2015; **Supplemental Figure 1G**). To determine the relative timing of WPR protein polarization, we examined SMCs in plants co-expressing either CFP-WPRA2 or RFP-WPRB2 with BRK1-CFP, PAN2-YFP, PAN1-YFP, or the F-actin marker FABD2-YFP **(Figure 3).** First, we compared the relative timing of the two WPR proteins (**Figure 3A**). In plants co-expressing CFP-WPRA2 and RFP-WPRB2, we observed 276 polarized SMCs, and in all cells CFP-WPRA2 and RFP-WPRB2 polarized at the GMC-SMC interface at the same time. We never observed polarization of one protein without the other. This is consistent with WPRA and WPRB proteins acting as a heterodimer. Based on this observation, we infer that polarization of either WPR protein at the GMC-SMC interface is likely indicative of the other.

**Figure 3.**
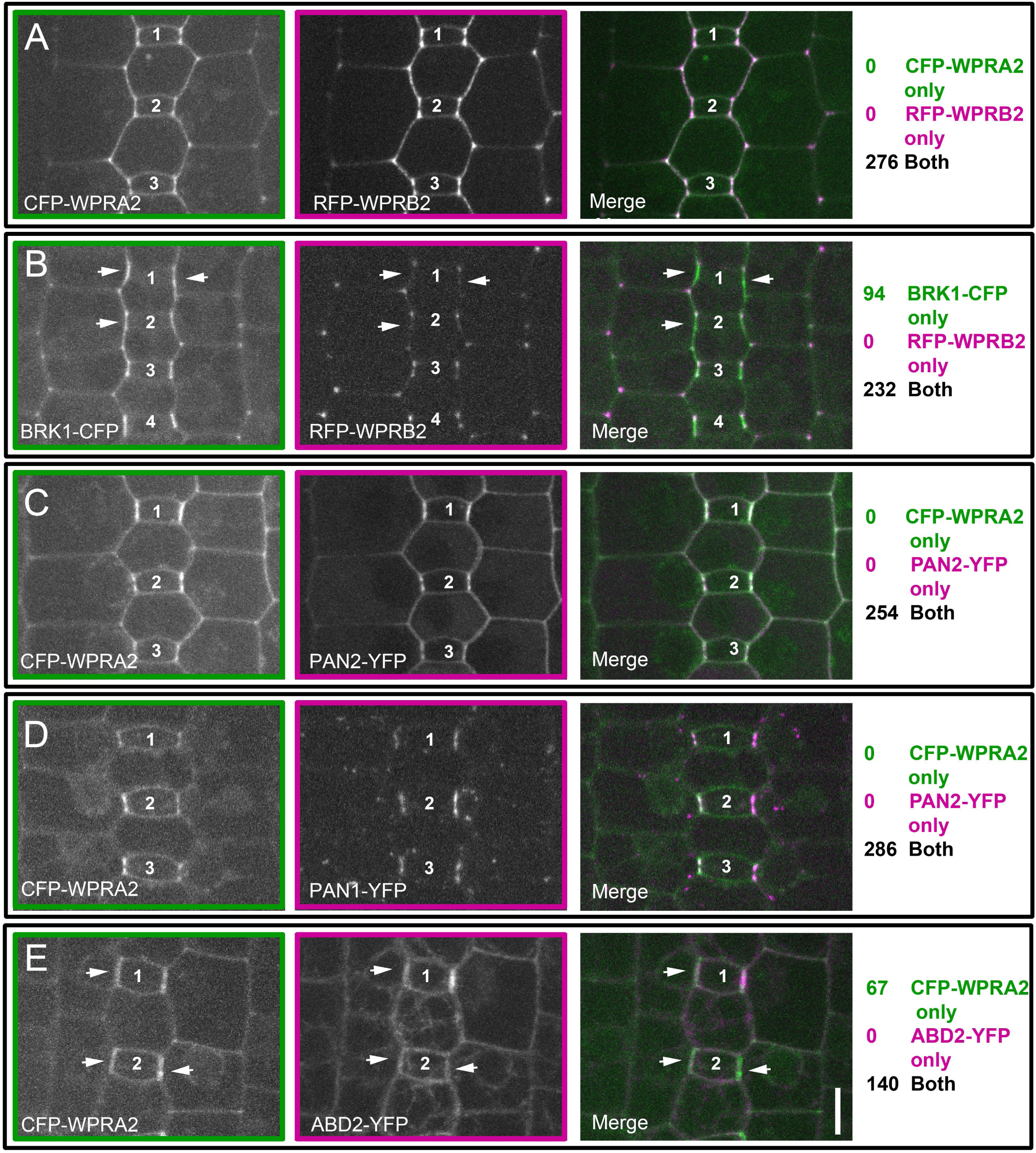
WPR proteins polarize in SMCs after BRK1 and before actin. The stomatal division zone of leaf 4 from plants co-expressing **(A)** CFP-WPRA2 & RFP-WPRB2; **(B)** BRK1-CFP & RFP-WPRB2; **(C)** CFP-WPRA2 & PAN1-YFP; **(D)** CFP-WPRA2 & PAN2-YFP; or **(E)** CFP-WPRA2 & FABD2-YFP were analyzed for enrichment at the GMC-SMC interface. Fluorescent proteins were scored as polarized when fluorescence was brighter at the SMC-GMC interface than at the cell periphery distal to the GMC. Within each panel, the same GMCs imaged in the different channels (and the subsequent merged panel) are numbered. Arrows point to SMCs where one fluorescent protein is polarized, but not the other. Counts of individual SMCs that had only one fluorescent protein, or both, are listed at right of the images. All images are at the same scale and are Z-projections of 7 confocal slices. Scale bar in the merged panel of E is 10 µm.

Next, we compared BRK1-CFP, the earliest known marker of polarity, and RFP-WPRB2. In plants co-expressing BRK1-CFP and RFP-WPRB2, we frequently saw that BRK1-CFP had become polarized while RFP-WPRB2 had not yet become polarized (94/232 cells). Arrows in **Figure 3B** indicate cells where BRK1 is polarized, but WPRB2 is not. Reciprocally, we never saw cells in which RFP-WPRB2 was polarized and BRK1-CFP was not polarized. This indicates that BRK1-CFP polarizes prior to RFP-WPRB2, and is consistent with previous findings that BRK1 is the earliest known marker of polarity **(Figure 3B)**. When CFP-WPRA2 was co-expressed with either PAN2-YFP **Figure 3C)** or PAN1-YFP **(Figure 3D)**, SMCs always showed co-polarization at the GMC contact site. The relative timing of WPR proteins and PAN1 receptors cannot be resolved from co-localization; our data indicates they polarizes at the same time. Accumulation of F-actin at the GMC-SMC interface is the last known marker of polarity. In SMCs co-expressing CFP-WPRA2 and FABD2-YFP, we could never see polarized accumulation of FABD2-YFP when CFP-WPRA2 was not polarized. However, we could see polarized accumulation of CFP-WRPA2 when actin had not yet accumulated in 67/140 cells, **(Figure 3E)**, suggesting WPRA2 polarizes prior to the actin patch in developing SMCs. Thus, WPR proteins polarize after BRK1-CFP, but prior to actin.

Co-expression of WPR proteins and PAN proteins indicate they polarize at (or very close to) the same time. To investigate the possible dependence of WPR polarization on the functions of known BRK-ROP-PAN pathway components, we examined the polarization of CFP-WPRA2 and RFP-WPRB2 in *brk1,pan1* and *pan2* mutants **(Figure 4)**. Homozygous mutant plants were compared to heterozygous siblings that were grown in parallel. Maize transgenics are generated in a hybrid background, and different inbred lines have a large amount of genotypic variation which may persist after backcrossing. By comparing mutants to phenotypically wildtype, heterozygous sibling plants, background genetic variation is accounted for, and any unexpected silencing of the transgene is also controlled for.

**Figure 4.**
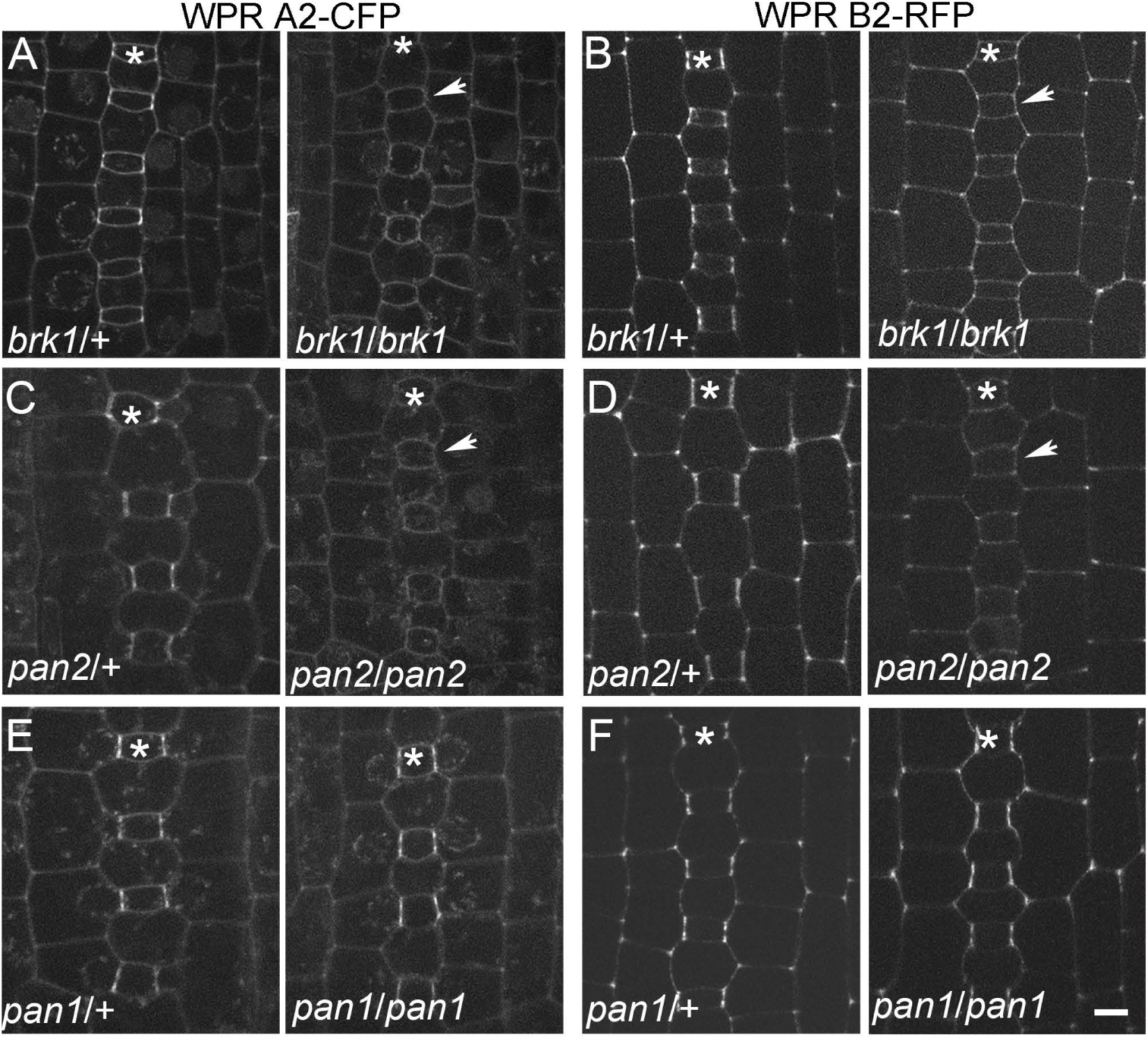
Polarization of WPR proteins depends on BRK1 and PAN2, but not PAN1. Localization of CFP-WPRA2 and RFP-WPRB2 in *brk1*, *pan2* and *pan1* mutants and non-mutant siblings was analyzed in developing leaf 4. CFP-WPRA2 **(A)** and RFP-WPRB2 **(B)** in developing stomata of *brk1* mutants and non-mutant (heterozygous) siblings. CFP-WPRA2 **(C)** and RFP-WPRB2 **(D)** in developing stomata of *pan2* mutants and non-mutant (heterozygous) siblings. CFP-WPRA2 **(E)** and RFP-WPRB2 **(F)** in developing stomata of *pan1* mutants and non-mutant (heterozygous) siblings. Asterisks mark GMC rows. Arrows point to SMCs without polarized localization of CFP-WPRA2 or RFP-WPRB2. 3-6 plants of each genotype were used for data collection. Scale bar, 5 µm.

Previously, it was shown that in *brk1* mutants, PAN1 and PAN2 do not polarize (Facette et al., 2015). Similarly, PAN1 does not polarize in a PAN2 mutant (Zhang et al., 2012). In *brk1* mutants, both CFP-WPRA2 and RFP-WPRB2 lost their polarized accumulation at the SMC-GMC interface (Figure 4A and 4B). This is consistent with the co-localization data, and indicates that WPRs act downstream of BRK1. Polarized localization of both CFP-WPRA2 and RFP-WPRB2 at the GMC contact site of SMCs is completely lost in *pan2* mutants **(Figure 4C and 4D).** In contrast, both WPR proteins remain polarized in *pan1* mutants **(Figure 4E and 4F)**. Thus, although we observed no timing difference in the polarized accumulation of PAN2, PAN1 and WPR proteins, PAN2 acts genetically upstream of WPRs and promotes their polarized accumulation. However, PAN1, which acts downstream of PAN2, is not required for WPR polarization.

### CRISPR/Cas9-Mediated Knockout of *Wprb1* and *Wprb2* Causes Subsidiary Cell Defects

To determine whether WPRs are required for asymmetric cell division during maize stomatal development, we used CRISPR/Cas9 to generate *wpr* mutants. The coding sequences of *Wpra1* and *Wpra2* are 92% identical, therefore both genes were targeted using the same two guide RNAs. *Wprb1* and *Wprb2* are 57% identical, and therefore two different guide RNAs were used in a single construct to separately target the two genes. Single mutants were identified for each gene **(Supplemental Figure 5A,B,C).** The edited lines were backcrossed with B73 and the transgene was segregated away. Single mutants were then crossed to obtain true-breeding double mutant lines. Despite PCR and sequence-based genotyping of many hundreds of plants from several different crosses, we never recovered *wpra1;wpra2* double homozygous mutants (**Supplemental Figure 5D**). This might reflect an essential shared function for the two genes in the male gametophyte, based on the high expression level of *wpra1* and *wpra2* in maize pollen (Supplemental Figure 5E). We were, however, able to generate *wprb1;wprb2* double homozygous mutants by crossing *wprb1/wprb1;wprb2/+* x *wprb1/+;wprb2/wprb2* plants (which generate double mutants at a higher frequency than a selfed double heterozygote). No stomatal phenotypes were observed in *wprb1* or *wprb2* single homozygotes, however *wprb1/wprb1;wprb2/wprb2* double mutants have an increased percentage of defective subsidiary cells relative to *wprb1/+;wprb2/+*, *wprb1/b1;wprb2/+ and wprb1/+;wprb2/b2* siblings (**Figure 5**). The abnormally shaped subsidiary cells are phenotypically similar to those found in *brk1, pan1* and *pan2* mutants (**Supplemental Figure 1H-K**; (Frank et al., 2003; Facette et al., 2015; Cartwright et al., 2009; Zhang et al., 2012). Stomatal density was unaffected in *wprb1/wprb1;wprb2/wprb2* mutants (**Supplemental Figure 6**). Since the aberrant subsidiary cell phenotype of *wprb1;wprb2* double mutants is mild relative to *pan1* or *pan2* mutants, we wanted to know if *wprb* mutants would enhance *pan2* phenotypes, similar to what has been observed in *rop2/4/9* mutants in combination with *pan1* (Humphries et al., 2011). However, we found that *wprb* mutations do not enhance the *pan2* phenotype (**Supplemental Figure 7**).We also examined *wprb1;wprb2* mutants for nuclear polarization and formation of the actin patch, two landmarks of polarity in SMCs (**Supplemental Figure 8**). No differences in nuclear polarization or actin patch formation were seen in *wprb1;wprb2* mutants. This may be because of the relatively mild defect or redundancy within the family. Higher order mutants, such as with the closely related *wprb3* mutant, could be informative.

**Figure 5.**
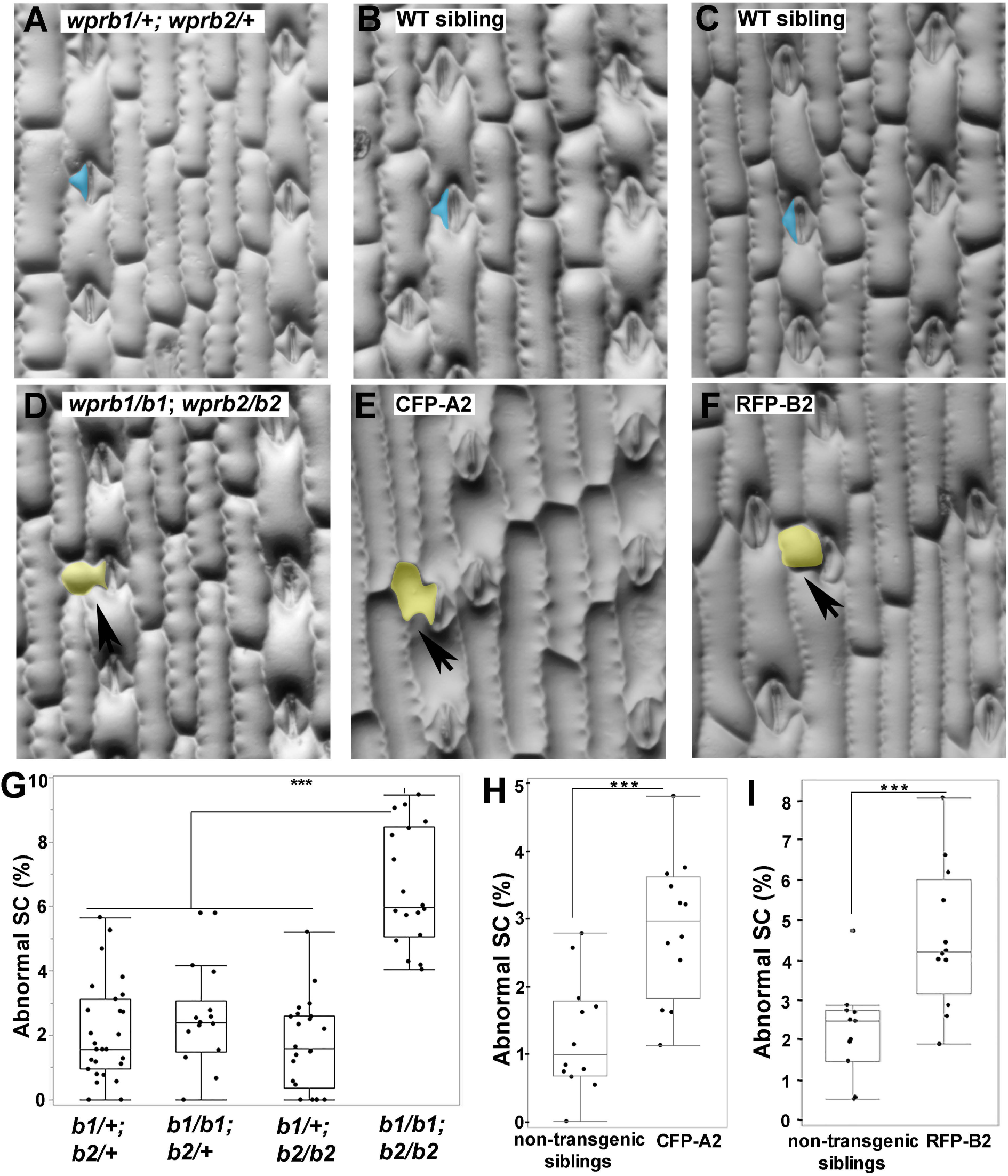
CRISPR-Cas9 induced *wprb1*;*wprb2* double mutants and CFP-WPRA2 and RFP-WPRB2 expressing lines have subsidiary cell defects. **(A-F)** Representative image of the third leaf epidermis of *wprb1/+; wprb2/+* and *wprb1/wprb1;wprb2/wprb2* mutants **(A and D)**, CFP-WPRA2 transgenic plants and non-transgenic siblings **(B and E)** and RFP-WPRB2 transgenic plants and non-transgenic siblings **(C and F).** Examples of normal subsidiary cells in A, B and D are false-colored blue, and abnormal subsidiary cells in D, E and F are false-colored yellow and marked with black arrows. (G) Quantification of abnormal subsidiary cells in *wprb1/+; wprb2/+* (n = 27 plants), *wprb1/b1;wprb2/+* (n = 14 plants), *wprb1/+;wprb2/wprb2* (n = 22 plants) and *wprb1/wprb1;wprb2/wprb2* (n=18 plants). For each plant 100-200 subsidiary cells were examined. p-value from Student’s T-test comparing each genotypes, ***P≤ 0.001. (H) and (I) Quantification of abnormal subsidiary cells in CFP-WPRA2 or RFP-WPRB2 expressing plants and non-transgenic siblings grown in parallel. (n = 11-12 plants and 120-200 cells per genotype). Student’s T-test were performed, ***p≤ 0.001.

In addition to loss-of-function phenotypes, we examined our native promoter-fluorescent protein fusion transgenics to determine if mild overexpression, resulting from co-expression of these FPs in a wild-type background with endogenous *Wpr* genes, would cause defective stomatal phenotypes. Plants expressing RFP-WPRB2 or CFP-WPRA2 in B73 indeed had mild subsidiary cell defects, further supporting that WPR proteins have a function in subsidiary cell division (**Figure 5 D, E**). These results indicate that both WPRA and WPRB family members participate in stomatal development, and suggest that WPRA1 and WPRA2 are important for the viability or function of male gametophytes.

### WPRB Proteins Directly Interact with F-actin

Since WPR proteins contain no discernable domains other than DUF827, their molecular role is unclear. We hypothesized that WPRs might interact with F-actin, based on the importance of actin in SMC polarization, and because *A. thaliana* WEB1 and PMI2 have roles related to actin-based regulated chloroplast movement (Kodama et al., 2010). Although we did not see obvious co-localization of WPR proteins with actin filaments in maize, membrane localization has been observed for other known actin-associated proteins such as BRK1 and WAL (Facette et al., 2015; Sugiyama et al., 2019). When overexpressed in tobacco cells, WAL localizes to actin cables (as opposed to membranes). To determine whether overexpressed WPRs similarly co-localize with F-actin *in vivo*, we transiently overexpressed GFP-tagged WPRA2, WPRB1, WPRB2 and WPRB3 proteins in tobacco epidermal cells. GFP-WPRA2 localized exclusively at the cell periphery/membrane **(Figure 6C)**. However, GFP-WPRB1 **(Figure 6D)**, GFP-WPRB2 **(Figure 6E)** and GFP-WPRB3 **(Figure 6F)** were observed in filament-like structures, and possibly also the plasma membrane. When GFP-WPRB2 is co-expressed with Lifeact-RFP (actin filament marker), the WPRB2-labeled filamentous structures extensively overlapped with Lifeact-RFP signals, particularly in the thick actin cables **(Figure 6L to 6N)**. We treated GFP-WPRB2 expressing tobacco leaves with Latrunculin B, an inhibitor of actin polymerization. The GFP-WPRB2 filamentous structures were disrupted, but remained intact in DMSO control **(Figure 6O and 6P, Supplemental Figure 9)**. Thus, WPRB2 is an F-actin associated protein.

**Figure 6.**
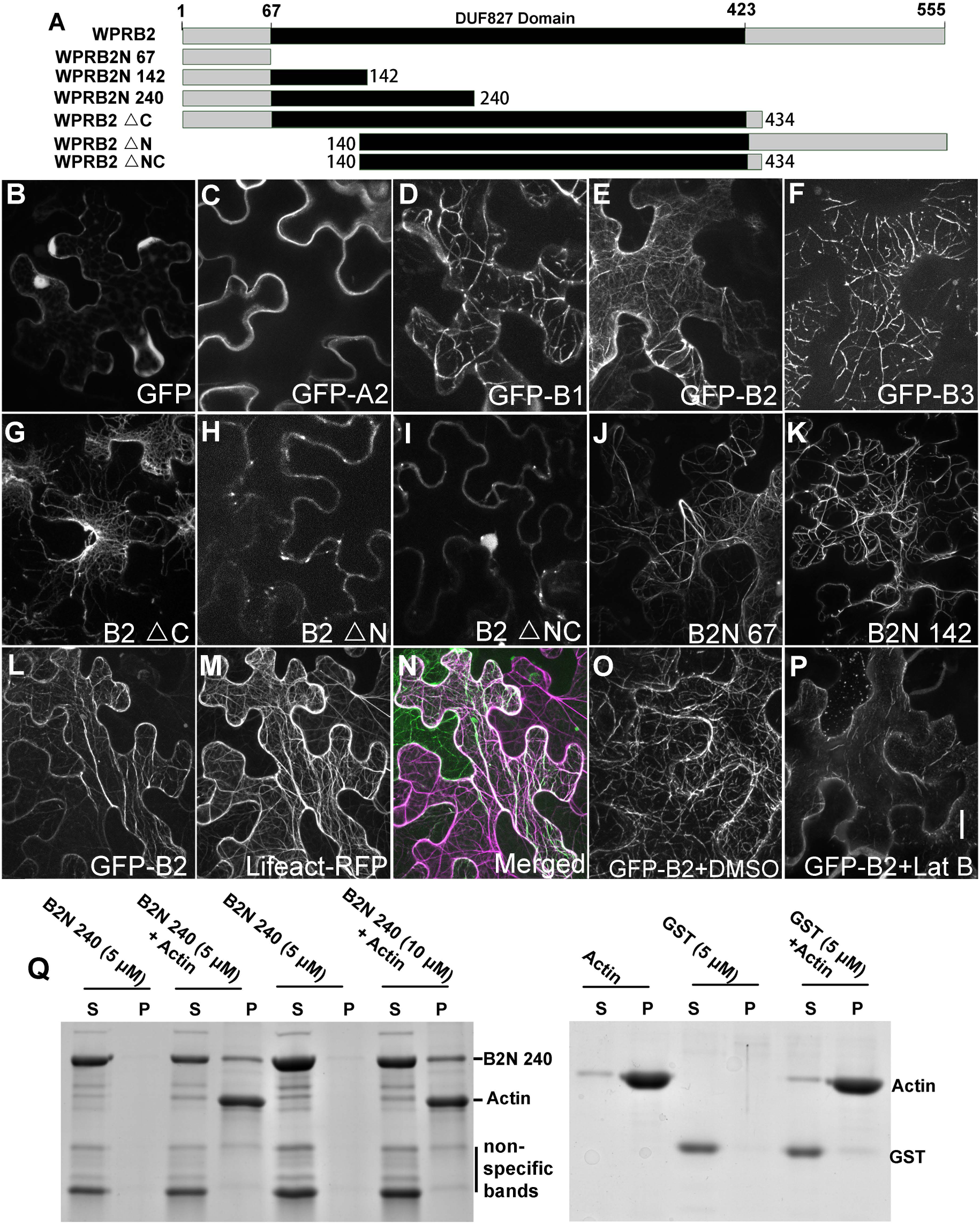
WPRB interacts with F-actin. **(A)** Schematic depicting WPRB2 and truncated versions. **(B-F)** Confocal images of transiently expressed full-length GFP fusion proteins in tobacco. Transient expression of GFP only **(B)**, GFP-WPRA2 **(C)**, GFP-WPRB1 **(D)**, GFP-WPRB2 **(E)** or GFP-WPRB3 **(F)**. **(G-K)** Transient expression of truncated WPRB2 proteins fused to GFP in tobacco leaves. Co-expression of GFP-WPRB2 **(L)** with Lifeact-RFP labeled actin microfilaments **(M)**; merged image **(N)** of the same cell. Tobacco leaves expressing GFP-WPRB2 were treated with DMSO (negative control, **O)** or 40 μM Latrunculin B **(P)** for 2 hours. All images at the same scale, Scale bar in **(P)** = 10 µm. **(Q)** High-speed co-sedimentation of GST-tagged WPRB2N240 with F-actin. After centrifugation at 100,000*g*, the proteins in the supernatant (S) and pellet (P) were resolved by SDS-PAGE and visualized with Coomassie Blue staining. GST protein was used as a negative control.

WPR proteins do not show significant sequence similarity to other known actin-associated proteins. To identify which region of the WPRB2 protein is responsible for its localization to filamentous structures, various WPRB2 fragments were used for transient tobacco expression assays **(Figure 6A)**. Surprisingly, the central conserved DUF827 domain is not responsible for filamentous localization. Instead, we found that the N-terminal 67 amino acids of WPRB2 is sufficient for filamentous localization in tobacco cells **(Figure 6J)**. Longer fragments containing the N-terminal 67 amino acids of WPRB2 were also found in filamentous structures, including GFP-WPRB2ΔC **(Figure 6G)** and GFP-WPRB2N142 **(Figure 6K)**. Any construct lacking the N-terminal domain does not label actin filaments, such as GFP-WPRB2ΔN **(Figure 6H)** and GFP-WPR B2ΔNC **(Figure 6I)**. We obtained similar results using full-length and truncated CFP-tagged constructs that were fused at C-terminus (rather than N-terminus) of WPRB2 **(Supplemental Figure 10)**. Conserved residues in the N-terminus of DUF827 proteins are highlighted in **Supplemental Figure 11.** Together, these data confirm that the N-terminal 67 amino acids of WPRB2 are necessary and sufficient for its filamentous localization.

To ask whether WPRB2 directly interacts with F-actin, we performed *in vitro* actin co-sedimentation assays. We used the slightly larger WPRB2N240 fragment, as the smaller WPRB2N67 fragment is similar in size to actin and made interpretation of the results difficult. WPRB2N240 is soluble in the absence of F-actin, but can be found in the pellet in the presence of F-actin **(Figure 6Q)**. The amount of WPRB2N240 appearing in the pellet is concentration dependent. Together, our localization and co-sedimentation experiments suggest that WPRB2 directly binds to F-actin via its N-terminal domain.

Physical interactions of WPR proteins with both PAN receptors and the actin cytoskeleton link together previously identified components of the BRK-PAN-ROP pathway. Potentially, WPRs may regulate actin polymerization, stability, or organization during some step of SMC polarization. Future studies will help resolve the relationship between WPRB proteins and actin. Intriguingly, during co-localization experiments in stably transformed maize plants described earlier (**Figure 3**), we noticed that the fluorescence intensity of the actin marker FABD2-YFP was reduced in RFP-WPRB2 co-expressing plants compared with RFP negative siblings, in all cell types. **Figure 7** shows the YFP channel in a plant expressing FABD2-YFP alone (**Figure 7Ai & Aii**), versus a sibling plant co-expressing both FABD2-YFP and WPRB2-RFP (**Figure 7Bi & Bii**), using identical acquisition settings. Fluorescence is barely visible in the co-expressing plants when the images are scaled at a range that is appropriate for the ABD2-YFP alone plants (Figure 7Ai vs.Bi). Reciprocally, when the YFP-channel is scaled appropriately for the co-expressing plants, the image is overexposed (Figure 7AIi vs.Bii).WPRB2-RFP intensity levels were similar in all sibling plants. We quantified the fluorescence in SMCs, and found decreased FABD2-YFP fluorescence in at the GMC contact site as well as at the other site was dramatically reduced, both at the polarized site as well as at the lateral side of the SMC (**Figure 7H**). The ratio of intensities indicates the polarized site is more affected (**Figure 7I**). This could be because of a critical function of WPRB2-RFP in promoting actin accumulation at the polarized site, rendering it more sensitive. It may be that more of an effect is seen at the polarized site, however, because more WPRB2-RFP accumulates there. Western blotting indicates that the overall actin levels are unchanged in RFP-WPRB2 expressing cells (**Supplemental Figure 12**). This suggests that the observed decrease in FABD2-YFP fluorescence could have a negative effects on actin stability, or alters actin in a way such that FABD2-YFP cannot bind as effectively.

**Figure 7.**
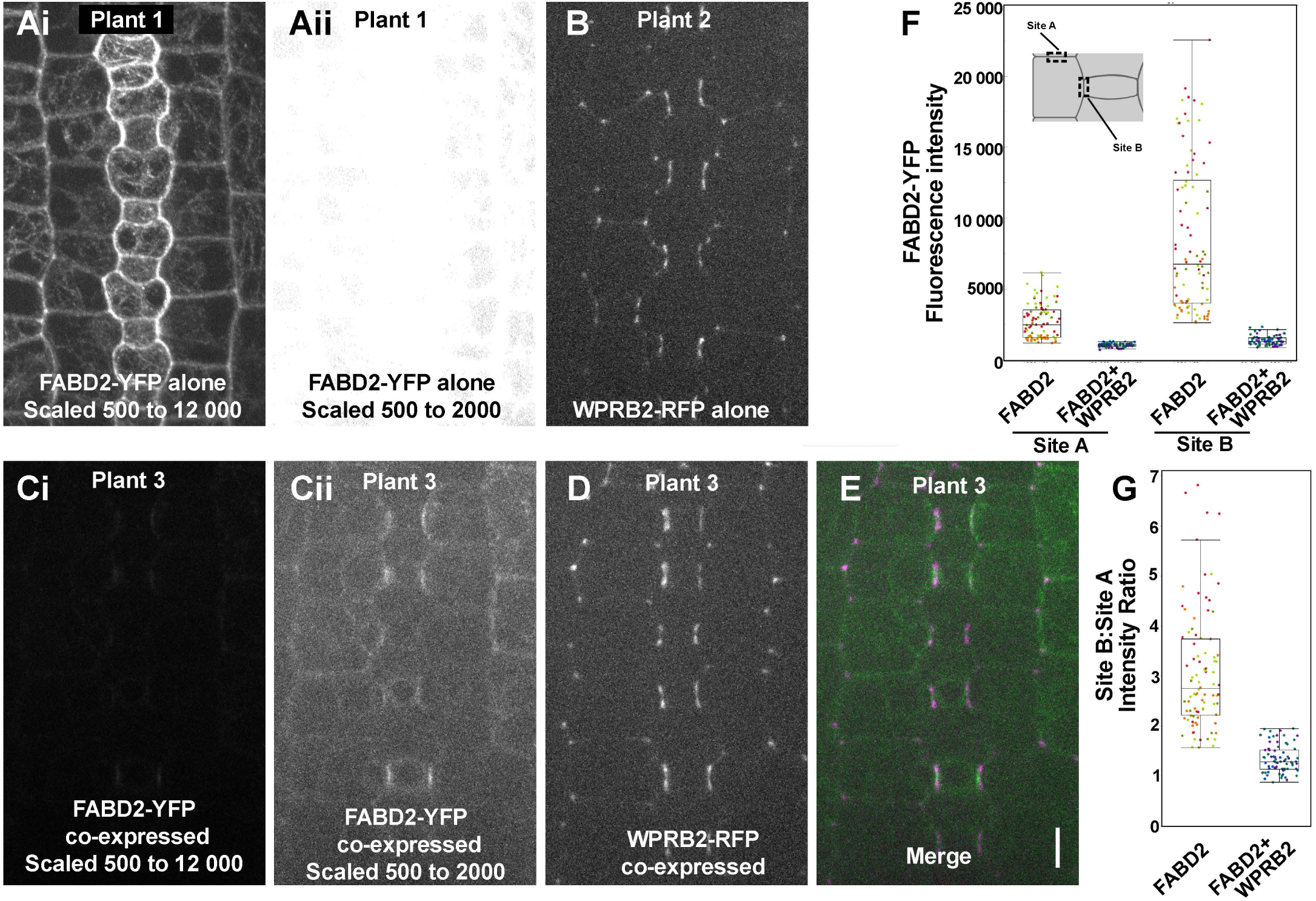
Fluorescence intensity of ABD2-YFP decreases in RFP-WPRB2 expressing plants. Plants expressing RFP-WPRB2 and FABD2-YFP were crossed, and the progeny independently segregated the two markers. (Ai & Aii) Plant only expressing FABD2-YFP. The identical image is shown in Ai and Aii. In Ai, the 16-bit image was scaled from 600 to 12000 prior to converting to 8 bit. Aii was scaled 500 to 2000. (B) Plant only expressing RFP-WPRB2. (C-E) Plant co-expressing FABD2-YFP (C, green in E) with RFP-WPRB2 (D, magenta in E). Ci and Cii show the identical image of the ABD2-YFP channel, where Ci is scaled the same as Ai and Cii is scaled the same as Aii, for comparative purposes. Images in B and D are scaled identically. Scale bar in E = 10 microns. (F) Quantification of fluorescence intensity of SMC lateral cell side (site A) and SMC-GMC interface (site B) in FABD2-YFP only plants (n = 5 plants, 292 cells) and RFP-WPRB2 and FABD2-YFP co-expressing plants (n = 3 plants, 164 cells). One sided t-tests indicate lower values in co-expressing plants (p < 0.0001). (G) Ratio of intensities measured in panel F. Two-sided test indicates a significantly lower ratio in the co-expressing cells (p<0.0001).

Our previous data indicates that expression of either CFP-WRPA2 or RFP-WPRB2 results in a low frequency of abnormal subsidiary cells (**Figure 5**), yet only WPRB (and not WPRA) proteins bind F-actin (**Figure 6**). We quantified whether CFP-WPRA2, like WPRB2, had an effect on FABD2-YFP intensity (**Supplemental Figure 13**). No similar decrease in FABD2-YFP intensity was observed in cells expressing CFP-WRPA2, consistent with the observation that only WPRB, and not WPRA, proteins bind F-actin.

Since actin is important in SMCs in early polarization, prior to WPR polarization (e.g., BRK proteins) and in late steps after WPR polarization (e.g. formation of the actin patch), we wanted to know if polarization of WPRB2 is maintained or stabilized by actin. We treated maize leaves expressing either RFP-WPRB2 or FABD2-YFP with Latrunculin B. After 4 hours, 40 µm Latrunculin B treatment led to efficient depolymerization of actin in SMCs, including the dense patches of actin at the SMC-GMC interface **(Supplemental Figure 14)**. However, disruption of actin does not affect the localization of RFP-WPRB2. This suggests that WPRB2 is not dependent on actin networks for maintaining its polarization. Together, these results suggest that WPRB2 interacts with actin filaments in a manner that de-stabilizes them, at least when present in excess of normal amounts. This offers a potential explanation for how WPRs may impact actin-dependent processes in polarizing SMCs.

## DISCUSSION

We demonstrated that WPR proteins function in the BRK-PAN-ROP pathway for subsidiary cell formation during maize stomatal development, and physically interact with the receptor proteins PAN2 and PAN1. Like other proteins in the BRK-PAN-ROP pathway, WPRs polarize in SMCs at the site of GMC contact. The polarized accumulation of WPRs in SMC depends on PAN2, and genetic evidence demonstrates a function for WPRs in promoting normal SMC divisions. Like WEB1 and PMI2 in *A. thaliana*, different WPR subfamily members form heterodimers, suggesting WPR subfamily members co-operate to fulfill their function in different cellular contexts (Kodama et al., 2010).

Our analyses indicate that WPR proteins act downstream of PAN2, but upstream of PAN1, yet physically interact with both PAN1 and PAN2. This is the first observed physical link between PAN2 and PAN1, and offers a hypothesis of how PAN2 may promote polarization of PAN1, i.e. indirectly via WPR recruitment. We predict that in *wpr* mutants, PAN1 will no longer become polarized; however gene redundancy within the WPR family makes this a non-trivial question to address. Future experiments with high-order *wpr* mutants and PAN1-YFP will further elucidate the function of WPRs in the pathway the pathway. Even though WPRA2 and WPRB2 no longer polarize in *pan2* mutants, they still localize to the plasma membrane, indicating that membrane localization is independent of PAN2.

Our data indicate that both WPRA and WPRB proteins localize predominantly to the plasma membrane, and may also have cytoplasmic localization. Similar to the BRK1, which is important for acting regulation, WPR proteins also localize to cell corners (Facette et al., 2015). Previous studies using transient protoplast expression of WEB1 and PMI2 indicate WEB1 localizes to the plasma membrane, and can recruit PMI2 there (Kodoma et al., 2010). An independent proteomic study indicated that WEB1 is predominantly localized to the cytoplasm but re-localizes to the membrane (and its phosphorylation status changes) upon blue light irradiation (Deng et al., 2014). No DUF827 family members have predicted transmembrane domains, indicating their membrane association is either peripheral or via interactions with membrane-localized proteins, or perhaps via membrane-associated actin networks. Besides WEB1 and PMI2, the other characterized DUF827 protein is TREPH1 (Wang et al., 2018). TREPH1 and maize WPRA2 are orthologues **(Supplemental Figure 2)**. A specific residue in the C-terminus of TREPH1 is phosphorylated in response to touch, and the phenotype of the *treph1* mutant is dependent on the phosphorylation status of this residue (Wang et al., 2018). However, the same residue in maize WPRA2 is not present. If all WPR proteins are similarly phosphoregulated, this suggests that other active, as of yet undiscovered kinases may be important for WPR regulation, since both PAN1 and PAN2 are pseudokinases.

Our analysis of CRISPR-Cas9 induced mutants of WPRs demonstrated that WPRs have diverse functions in maize. We were unable to recover *wpra1*/*wpra1*;*wpra2*/*wpra2* double mutants. WPRA1 and WPRA2 are 89% similar in protein sequence, and are highly expressed in pollen. CFP-WPRA2 polarly localizes to the subapex of maize pollen tubes **(Supplemental Figure 5)**, raising the possibility that WPRA proteins play a role in pollen function. Additionally, we found *wprb1;wprb2* double mutants affect subsidiary cell formation, albeit not to the same extent as *pan* mutants. This may be caused by functional redundancy with other *Wpr* genes, such as *Wprb3*.

Our discovery that WPRB proteins directly bind actin is significant as it physically links them to actin network function. WPR proteins do not share any sequence homology to known actin binding proteins, however several DUF827 proteins were predicted, based on structural homology, to potentially bind the cytoskeleton (Gardiner et al., 2011). Surprisingly, the central conserved DUF827 domain is not the actin-binding domain. We mapped a 67 amino acid sequence that is necessary and sufficient for actin-binding, thereby identifying a putative novel actin binding domain. Within this sequence, there appears to be a motif that is conserved in many, but not all DUF827 proteins **(Supplemental Figure 8)**. Even though WPRA2 also contains most of this conserved domain, we did not observe co-localization of WPRA2 to the actin cytoskeleton. This could be because the transient expression does not reflect *in vivo* function. Alternatively, small differences between WPRA2 vs. B2 sequence may account for its lack of actin binding (e.g, one of a pair of glycines conserved in many WPR proteins is aspartate in WPRA2). Further refining the region of interaction, and determining if other DUF827 proteins bind actin, would inform their function.

The significance of WPR interaction with the actin cytoskeleton is unknown, and is complicated by the fact that there are several actin-related processes that occur during SMC polarization. The most plausible are: a function related to the SCAR/WAVE complex that includes BRK1, which polarizes and functions very early to promote SMC polarization; a role in localization of polarity determinants (such as vesicle trafficking); a role related to actin patch formation or function; or a role related to actin-based nuclear migration. These possibilities are not mutually exclusive. Actin filaments can facilitate polarized protein localization. For example, polarization of PIN requires an intact actin cytoskeleton (Kleine et al., 2008). It is plausible that actin networks are responsible for the polarization of PAN1 and/or PAN2, or even the WPR proteins themselves. This function would be distinct from that of the actin patch, which appears long after PAN2, PAN1, or WPR proteins polarize. When actin patches were disrupted with Latrunculin B, this did not result in the loss of already polarized RFP-WPRB2 in maize leaves, suggesting the actin patch does not influence WPR protein localization (and possibly function). A clue to what WPR proteins could be doing *in vivo* can be inferred from our observation that RFP-WPRB2 transgenic plants WPRB displayed less F-actin than controls. This suggests WPRB2 negatively affects the stability of actin filaments, and might promote F-actin depolymerization. Alternatively, WPRB proteins could compete with ABD2 for binding sites on the F-actin polymer, or somehow modify F-actin such that ABD2 does not bind as effectively. Future *in vitro* and *in vivo* work characterizing the molecular role of WPR proteins on actin dynamics, including the nucleation, severing and bundling activities will help determine the function of this gene family – not only in cell polarization, but also during mechanosensing (i.e., TREPH1 function) or organelle movements (i.e. WEB1 and PMI2 function).

## METHODS

### Plant Materials and Plant Growth Condition

BRK1-CFP, PAN2-YFP, PAN1-YFP, and YFP-ABD2-YFP stable transgenic lines, and *brk1*, *pan1* and *pan2* mutants were described previously (Cartwright et al., 2009, Zhang et al., 2012; Facette et al., 2015). Plants used for analysis were grown for 10d to 14d in a greenhouse maintained between 20°C and 29°C. Plants were grown in Pro-mix Professional soil supplemented with . Supplemental LED lights (Fluence VYPR series) were used to maintain a 16-hour day length.

### Generation of transgenic maize lines

Transgenic lines expressing CFP-WPRA2 or RFP-WPRB2 were created using a genomic construct. Primers are listed in **Supplemental Table 2.** An approximately 3kb sequence upstream of the start codon (promoter and 5’ UTR) was amplified from B73 genomic DNA with primers WPRA2-F & WPRA2-P2 or WPRB2-F & WPRB2-P2. Genomic DNA, including all exons and introns 1kb after the stop codon (3’ UTR), was amplified from B73 genomic DNA using primers WPRA2-P3 & WPRA2-R, or WPRB2-P3 & WPRB2-R. CFP or RFP was amplified with primer Tag linker-F and Tag linker-R as described previously (Mohanty et al., 2009). N-terminal fluorescent protein fusions were assembled through fusion PCR and cloned into pDONR221. Gateway LR reactions (Thermo Fisher) were used to recombine the assembled fusion protein into the binary vector pAM1006 (Monhanty et al., 2009). The constructions were verified and introduced into Hi-II maize via *Agrobacterium tumefaciens*–mediated transformation by Wisconsin Crop Innovation Center plant transformation facility. Primary transformants were crossed to B73 to produce T1 progeny used for immunoprecipitation experiments, and T1s were crossed with BRK1-CFP, PAN2-YFP, PAN1-YFP, and YFP-ABD2-YFP to produce T2 progeny used for the imaging experiments. Primary transformants were also crossed to *brk1*, *pan1* and *pan2* mutants, and the T1 progeny backcrossed again to mutants, to produce homozygous *brk1*, *pan1* and *pan2* mutants expressing CFP-WPRA2 and RFP-WPRB2. In all cases, sibling plants grown simultaneously were used as controls.

### Yeast Two-Hybrid Analysis

The GAL4-based yeast two-hybrid interaction system was used for interaction assays. The cDNA fragments encoding intracellular portions of PAN1 and PAN2 were cloned into pASGW-attR plasmid as a bait via Gateway cloning as described previously (Zhang et al., 2012). cDNA was prepared from the maize leaf stomatal division zone. Full length cDNAs of *Wpra1, Wpra2, Wprb1, Wprb2* and *Wprb3* cDNA were amplified from a maize leaf division zone cDNA preparation with primers WPRA1topo1-F + WPRA1topo-R (*Wpra1*), WPRA2topo1-F + WPRA2topo-R (*Wpra2*), WPRB1topo1-F + WPRB1topo-R(*Wprb1*) and WPRB2topo1-F + WPRB2topo-R (*Wprb2*) (**Supplemental Table 1**). These fragments were cloned into the entry vector pENTR/D-TOPO (Invitrogen) and further transferred destination vectors pASGW-attR (bait) and pACTGW-attR (prey) (Nakayama et al., 2002). Bait and prey plasmids were co-transformed into AH109 yeast strain. Yeast was grown on medium selecting for diploid cells (SD -Leu-Trp) or detecting interactions (SD -Leu-Trp-Ade-His) for two days at 30°C.

### Confocal Microscopy

Plant tissues were observed using a custom-built spinning disc confocal unit (3i) equipped with an inverted fluorescence microscope (IX83-ZDC, Olympus) CSU-W1 spinning disc with 50 micron pinholes (Yokogawa), a Mesa Illumination Enhancement 7 Field Flattening unit (3i), an Andor iXon Life 888 EMCCD camera and a UPLANSAPO ×60 Silicone Oil-immersion objective (NA = 1.20, Olympus) and 4 laser stack with TTL controller (3i). For CFP, YFP and RFP imaging of transgenic maize plants, a 445/515/561 dichroic (Chroma) was used. All emission filters are from Semrock. A 445 nm laser line and 483/45 emission filter (CFP), 514 nm laser and 542/27 emission filter (YFP) or 568 laser with 618/50 emission filter (RFP) was used. For dual imaging of GFP and RFP (tobacco expression) a 405/488/561/640 dichoric (Chroma) was used, with a 488 nm laser and 525/30 emission filter (GFP) and/or 568 laser with a 618/50 emission filter. Image processing was performed using Image Fiji and Adobe Photoshop version 8.0 using only linear adjustments and preserving hard edges.

Quantification of ABD2-YFP fluorescence intensity was performed using FIJI. Max projections of 7 slices of 16-bit images taken on the same day with the same acquisition settings were used. Sibling plants were used as controls.

### Transient expression in *N. benthamiana*

To generate 35S-expressed N-terminal GFP-tagged constructs (and truncated derivatives) and C-terminal CFP-tagged constructs (and truncated derivatives) for transient tobacco expression, coding sequence were PCR-amplified from maize leaf cDNA with the appropriate primers (**Supplementary Table 1**) and cloned into the pENTR/D-TOPO entry vector (Thermo Scientific). Clones were recombined with the vectors pSITE-2CA and pSITE- 1NB using LR Clonase Mix II (Thermo Scientific) to form N-terminal GFP or C-terminal CFP (Romit Chakrabarty, 2006).

For transient expression in *N. benthamiana*, *Agrobacterium* strain GV3101 harboring different constructs were resuspended in infiltration buffer (10 mM MES (pH 5.7), 10 mM MgCl_2_, 50 mg/L acetosyringone), and adjusted OD600 to 1.0. Equal volumes of cultures containing different constructs were mixed for co-infiltration. The resulting cultures were infiltrated into leaves of 3-to 4-week-old N. *benthamiana* plants. Leaf samples were harvested 48 hours after infiltration.

### Generation and Purification of WPRA1/A2 and WPRB2-Specific Antibody

Peptides corresponding to amino acids 267 to 288 of maize WPRA1 and WPRA2 (LRNDFDPAAYDSLKEKLEQTNS), and amino acids 446-465 of WPRB2 (HPAPRSRDSQNMDIVGVSKGC) were synthesized, conjugated to KLH, co-injected into rabbits and used for polyclonal antibody production in rabbits by Pacific Immunology. And the resulting sera was affinity purified against the corresponding peptide using columns provided by Pacific Immunology. Affinity purified antibodies were eluted using Gentle Ag/Ab buffer (Pierce), desalted against TBS Buffer, and concentrated to 1 mg/ml using a 30K MWCO concentrator (Pierce).

### Maize Division Zone Membrane Protein Extraction, Western Blotting, and Co-IP/MS Analysis

Protein extraction for western blotting of WPR proteins and co-IPs were performed as described previously (Facette et al., 2015). Briefly, 10-14 day old plants were used to isolate the basal 0.5-2.5 cm of unexpanded leaves 4-6, the leaf bases were ground in liquid nitrogen. Membrane fractions of extracts from these tissues were prepared. For western blotting, ∼0.5 grams (3-5 plants) of tissue was used and for mass spectrometry, 1.5 grams (8-10 plants) were used. per 0.25 g of grounded tissue was mixed with 1ml extraction buffer (50mM Tris, pH 7.5, 150 mM NaCl, 5mM EGTA, 5mM EDTA 0.3% β-mercaptoethanol, 1% Sigma Plant Protease Inhibitor), the mixture was homogenized for 10s using an Omnitip homogenizer on ice. All subsequent steps were conducted at 4°C. The mixture was centrifuged twice at 13000*xg*. The supernatant was pelleted at 110,000xg for 45 min. Solubilization buffer (50 mM Tris pH 7.5, 150 mM NaCl, 1% NP-40, 10% glycerol, 0.05% Sodium deoxycholate) was then added to the pellet and following sonicated 3×15s and left rotating 1 hour. For western blotting, extracts from B73 plants and corresponded mutants were separated via SDS-PAGE and analyzed using anti-WPRA1/A2 at 1 μg/ml.

For western blots to detect actin in B73 and RFP-WPRB2 transgenic plants, the basal 0.5-2.5 cm of unexpanded leaves 4-6 from 14 days plant were ground in liquid nitrogen. Groundtissue was mixed with 500 µL extraction buffer (50mM Tris, pH 7.5, 150 mM NaCl, 5mM EGTA, 5mM EDTA 0.3% β-mercaptoethanol, 1% Sigma Plant Protease Inhibitor) and homogenize immediately by Omnitip homogenizer for 10s on ice. Then, the mixture was centrifuged at 15,000 x g for 10 min at 4 °C. Transfer supernatant (protein extract) to a new tube and keep on ice for immediate use. 100 ng extracted protein for each plants were separated via SDS-PAGE and analyzed using anti-actin (Sigma A0480, MFCD00145889) at 1:1000.

Co-IP/MS experiments were performed using the solubilized membrane proteins. Immunoprecipitation of PAN2-YFP was performed as in Facette et al. (2015) using anti-GFP beads (Miltenyi). PAN2-YFP immunoprecipitations and subsequent mass spectrometry runs were performed in parallel with B73 (control), BRK1-CFP, PAN1-YFP, PDI-YFP, PIN1-YFP and RAB11D-YFP using anti-GFP beads (Miltenyi). Note that these co-immunoprecipitations, except PAN2-YFP and PAN1-YFP, were presented previously in Facette et al. (2015) A WD-score (Sowa et al., 2009) was calculated to determine high confidence interactors. Tryptic digestion and mass spectrometry was performed using a Q-Exactive mass spectrometer (Thermo Scientific, San Jose, CA) as described (Facette et al., 2015).

Anti-WPRA1/A2 and anti-RFP (Rabbit polyclonal, Invitrogen R10367, Lot 2127438) were coupled with Dynabeads prepared according to the Dynabeads kit instructions (Thermo Fisher). The coupled WPRA1/A2 antibody-beads complex was added to the membrane protein extracts from B73 plants; uncoupled beads were used as a negative control. Anti-RFP coupled beads were added to the membrane protein extracts from RFP-WPRB2 transgenic plants, and B73 plants as a negative control. All these samples were incubated rotating at room temperature for 30 min. Then the Dynabeads-CO-IP complex was washed according to Dynabeads kit instructions. After washing, the Dynabeads-CO-IP complex was digested with trypsin and run with the mass spectrometer.

For Co-IP using anti-GFP to pull down WPRA2, anti-GFP beads (Miltenyi) were added to the extracts from CFP-WPRA2 transgenic plants and B73 plants (negative control), then the extracts were rotated for an additional 30 minutes. Miltenyi µ columns were equilibrated using a solubilization buffer as described by instruction. The extracts were applied to the columns, the columns were washed and eluted as Miltenyi µ columns kit instruction, the eluted samples were digested with trypsin and run with the mass spectrometer.

For all WPR immunoprecipitations, peptides were separated by reverse-phase chromatography using nano-flow EASY-nLC 1000 UHPLC coupled to Orbitrap Fusion mass spectrometer (Thermo Scientific) with PepMap RSLCnano column (75 μm ID, 15 cm). Peptides were eluted over a 90 minute 5-35% ACN gradient at 300 nL/min. Survey scans were measured in the Orbitrap analyzer at 60,000 resolution. Data-dependent MS/MS data were collected in the linear ion trap using a 2-second cycle time with a full MS mass range from 400-1800 m/z. Peptides (charge state 2-6) were fragmented using higher energy collisional dissociation using a normalized collision energy setting of 27. A dynamic exclusion time of 5 seconds was used and the peptide match setting was enabled. RAW files were analyzed in Proteome Discoverer 2.4 (Thermo Scientific) using the SEQUEST search algorithm using version 4 of the maize genome (B73 RefGen_v4). The search parameters used are as follows: 10 ppm precursor ion tolerance and 0.4 Da fragment ion tolerance; up to two missed cleavages were allowed; dynamic modifications of methionine oxidation and N-terminal acetylation. Peptide matches were filtered to a protein false discovery rate of 5% using Percolator algorithm. Peptides were assembled into proteins using maximum parsimony and only unique and razor peptides were retained for subsequent analysis.

### Genomic editing of maize WPRs by CRISPR–Cas9

A dual gRNA maize CRISPR/Cas9 system was used to generate *Wpr* mutants (Char et al., 2017). Transgenic Hi-II maize plants were generated at the Iowa State Plant Transformation facility, expressing Cas9 from *Streptococcus pyogenes* under the maize UBI1 promotor and two guide RNAs under the rice U6.1 and U6.2 promotor, respectively. To create *wpra1* and *wpra2* mutants, two tandem gRNAs, gRNA1: GAAACATCTTTGGATATG and gRNA2: GTGCAAGCACACGAAGAAG were used to target identical sequences within *Wpra1* and *Wpra*2. To create *wprb1* and *wprb2* mutants, guide RNAs targeting *Wprb1* using gRNA1: GCTACATGTGATCTGGCTG or *Wprb2* using gRNA2: GCCTCCGTCGAGTCGCTG were used. Genomic edits were verified by Sanger sequencing of the target regions. Plants harboring edits were crossed to B73. Edited plants that no longer contain the transgene were confirmed and used for further analyses.

### Phalloidin staining

Phalloidin staining in maize leaves was performed as previously described (Cartwright et al., 2009, Nan et al., 2019). The basal 0.5-2.5 cm of leaf 4 from *wprb1/+;wprb2/+* and *wprb1/b1;wprb2/b2* mutants were fixed and stained with Alexa fluor 488-phalloidin (Thermo Fisher). Nuclei and cell walls were stained using 10 μg mL-1 propidium iodide (Thermo Fisher). Samples were mounted in ProLong Gold Antifade (Thermo Fisher).

### Immunofluorescent detection of WPR proteins

Immunolocalization of WPRA2 and WPRB2 were performed in leaf tissue excised from the basal 1 to 3 cm of unexpanded leaves of 2-week-old plants as described previously (Cartwright et al., 2009, Nan et al., 2019) using affinity-purified anti-WPRA1/A2 or WPRB2 at 2 µg/ml. Secondary Alexa Flour 488-conjugated anti-rabbit (Invitrogen) were used at a dilution of 1:500. Nuclei were stained with 10 µg/mL propidium iodide (Sigma-Aldrich) prior to mounting in in ProLong Gold Antifade (Thermo Fisher).

### Bacterial Expression of WPR proteins

The DNA sequence of WPRB2N (1-240), which was used in the actin co-sedimentation assay, was amplified by PCR using the primers listed in Supplemental Table 2 and cloned into pGEX4T-1. Fragments of WPRA1 (63-503) WPRA2 (63-570) WPRB1 (73-421) and WPRB2 (78-429), which were used to confirm the specificity of the WPRA1/WPRA2 antibody, were also amplified by PCR using the primers listed in Supplemental Table 2 and cloned into pGEX4T-1 (Novagen) using BamHI/EcoRI sites to N-terminal glutathione S-transferase (GST) tag fusions. These constructions were transformed into *Escherichia coli* BL21 host strains (Novagen). GST fused proteins were induced with 0.5 mM isopropyl β-d-1-thiogalactopyranoside for 20 hours at 28°C. Affinity purified using glutathione-Sepharose 4B column (GE Healthcare) was carried out as described (Harper and Speicher, 2011).

### F-Actin Co-sedimentation Assay

WPRB2N240 was dialyzed overnight against buffer A3 (10 mM Tris-HCl, 0.2 mM CaCl2, 0.2 mM ATP, and 0.5 mM DTT, PH=7.0). Prior to use, the protein was further purified by centrifugation at 100,000g for 1 hour at 4°C and only soluble protein was used for the assay. A high-speed co-sedimentation assay was performed as described previously (Xiang et al., 2007), Briefly, 5 μM and 10 μM WPRB2N240 alone or mixed with 4 μM preformed F-actin and incubated in 200 μL volume of buffer A3 for 1 hour at 25°C. The samples were centrifuged at 100,000g for 1 hour at 4°C. GST protein was used as a negative control. The supernatants and pellets were separated and subjected to SDS-PAGE and visualized by Coomassie Blue staining.

### Latrunculin B treatment

To transiently depolymerize actin microfilaments in maize leaves, leaf 4 of two-week-old plants were excised and the basal 1 to 3 cm part was immersed in 40 μM Lat B (20 mM stock in DMSO) for 4 hours at room temperature. To examine the LatB effect on the localization of GFP-WPRB2 in tobacco leaves, the GFP expressed tobacco leaves were cut into 0.5 cm×0.5 cm and covered with 40 μM LatB for 2 hours at room temperature.

### Protein tree construction

Amino acid sequences of maize WPR homologs were identified using pBLAST against maize genomes. The protein tree was constructed in Molecular Evolutionary Genetics Analysis 5 (MEGA 5) software (Tamura et al., 2011) with the following parameters: multiple amino acids sequence alignment with ClustalW, phylogenetic construction with the maximum likelihood method, and bootstrap tests of 1,000 replicates. WPRs sequences used for the phylogenetic analysis are provided in **Supplementary File 1**.

### Statistical Tests

T-tests and one-way ANOVA test were performed in JMP (SAS). Chi-squared test was performed using an online calculator released by GraphPad Prism (https://www.graphpad.com/quickcalcs/chisquared1/) with expected values calculated by hand.

### Accession Numbers

Gene and protein sequences can be found at MaizeGDB www.maizegdb.org. Accession numbers for Version 4.0/Version 5.0 of the B73 genome are: *Wpra1:* Zm00001d023629/Zm00001eb408590; *Wpra2*: Zm00001d041088/ Zm00001eb133280; *Wprb1:* Zm00001d0475516/Zm00001eb395070; *Wprb2:* Zm00001d007164/Zm00001eb111490; *Wprb3:* and Zm00001d010610/ Zm00001eb351980; *Brk1:* no v4.0/ Zm00001eb259430; *Pan2:* Zm00001d007862/Zm00001eb117610; *Pan1*: Zm00001d031437/ Zm00001eb034900.

## ACKNOWLEDGEMENTS

We thank Steve Eyles at the UMass Institute of Applied Life Sciences Mass Spectrometry Facility, RRID:SCR_019063. We thank Chris Phillips, Bob Skalbite, Neal Woodward, and other greenhouse and farm staff at the University of Massachusetts Amherst for essential help. This work was supported by a grant from NSF (IOS-1754665) to MRF and NIH (DP2-GM119132) to EJB.

## AUTHOR CONTRIBUTIONS

M.F. and Q.N. designed and performed the research, analyzed the data and wrote the manuscript. B.Y. (UCSD) and E.J.B. performed the mass spectrometry for the initial set of co-IPs. S.N.C. and B.Y.(MO) designed the WPRs CRISPR sgRNAs and prepared the constructs for maize transformation.

## Supplemental Data

**Supplemental Table 1.**
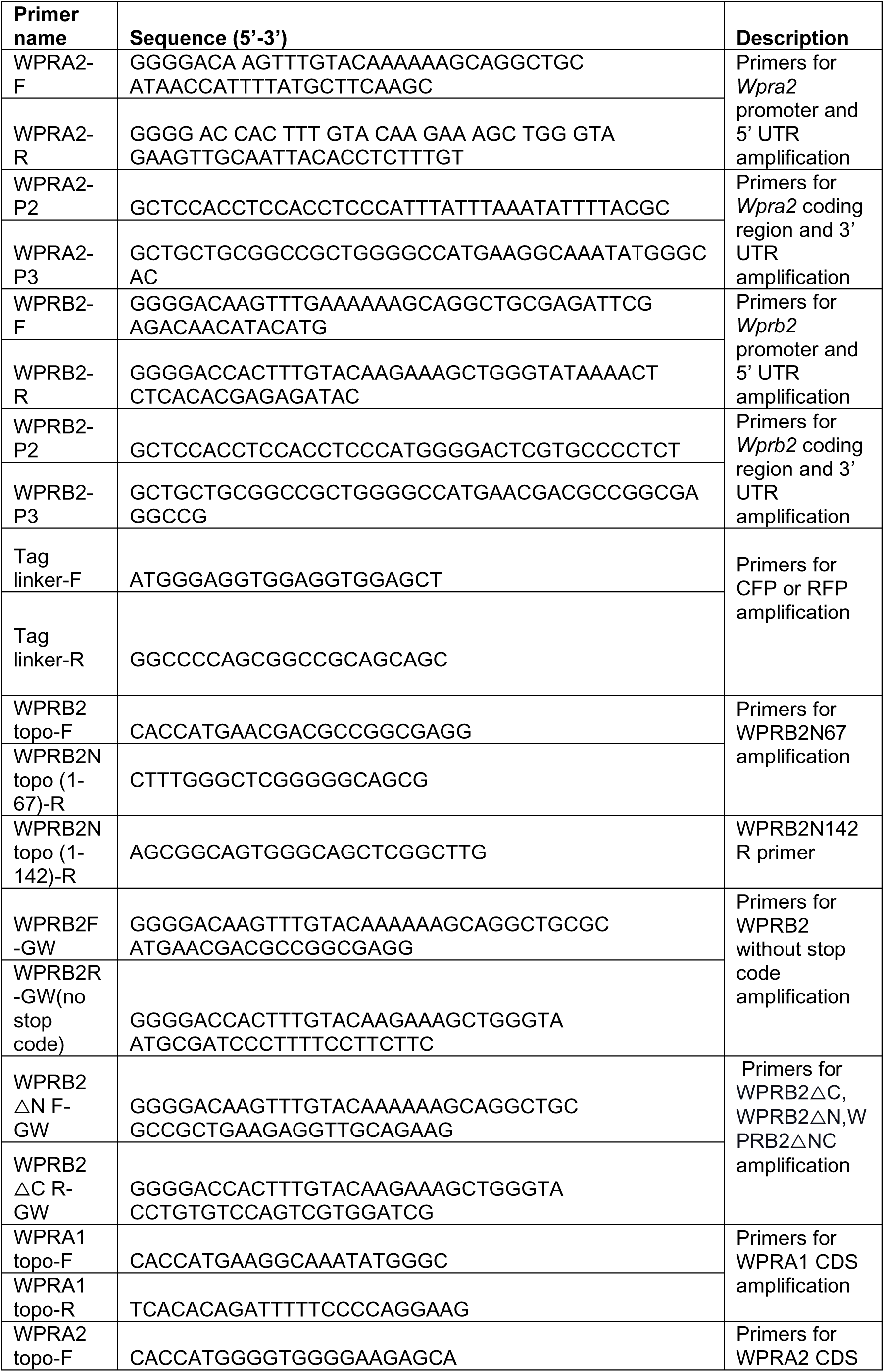
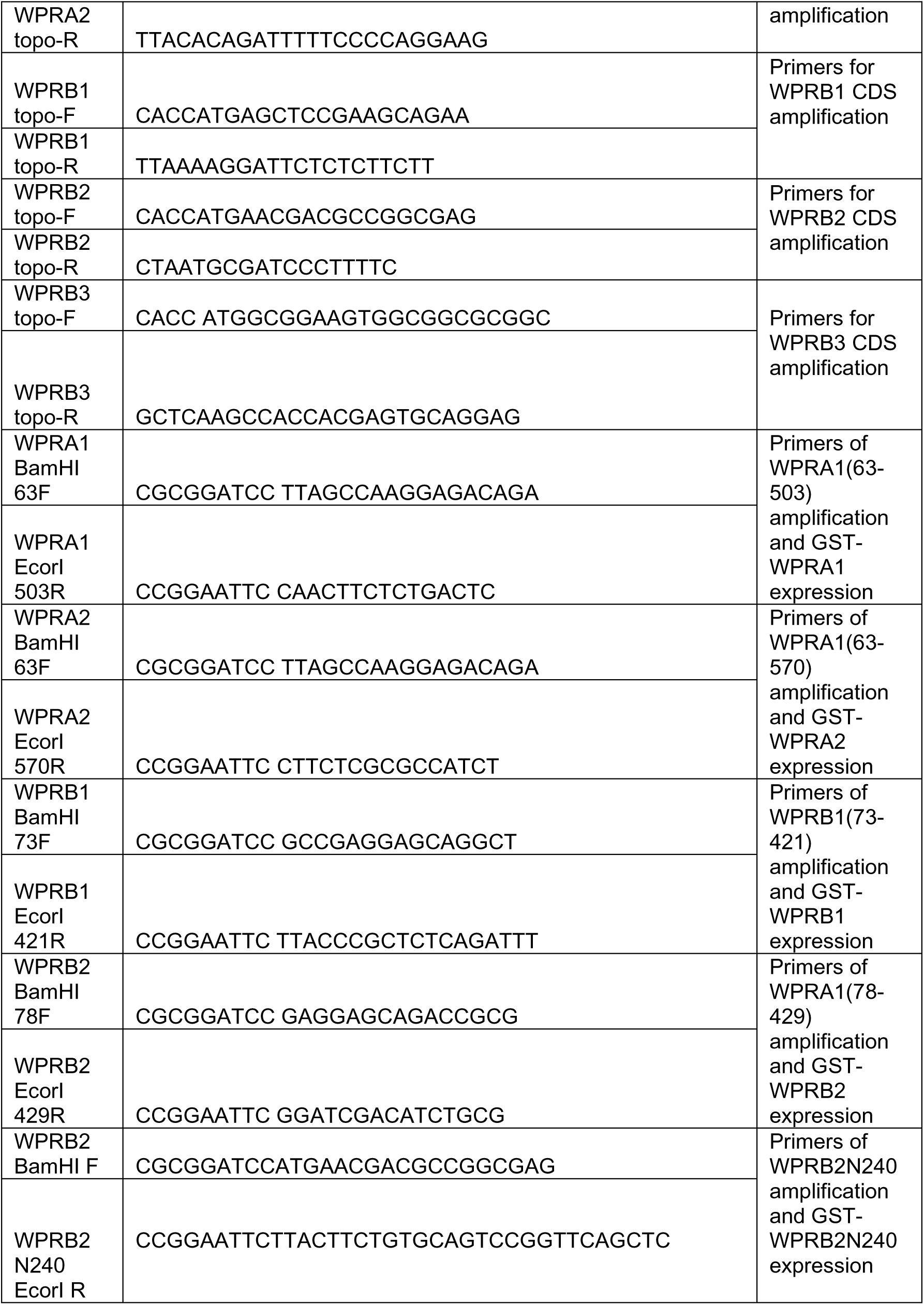
Primer sequences used in this study.

**Supplemental Figure 1.**
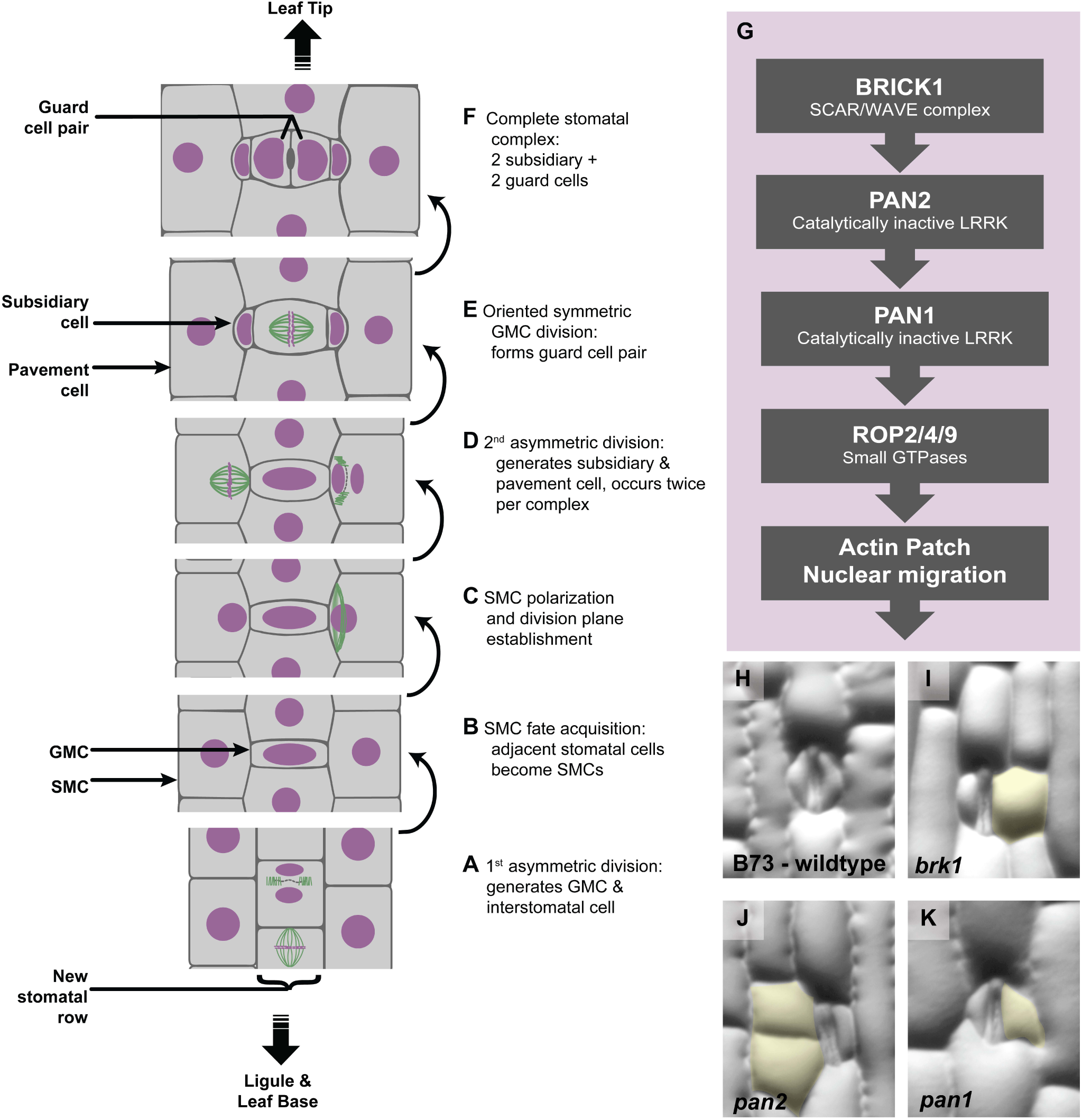
Progression of maize stomatal development in leaves. Unexpanded maize leaves mature in a gradient with the developmentally youngest cells towards the base and the developmentally oldest cells towards the tip. **(A)** Stomatal complex formation is initiated by a transverse asymmetric division in a new stomatal row. **(B)** Pavement cells flanking the GMC acquire a SMC fate. At this stage, the nuclei are not yet polarized, but proteins (not shown) in the SMC may be polarized towards the GMC. **(C)** SMC nuclei polarize towards the GMC; preprophase band forms. **(D)** The SMCs divide, with each division forming **(E)** a subsidiary cell and a pavement cell. The GMC undergoes an oriented, symmetric division symmetrically to form (**F**), the guard cell pair. Each stomatal complex is made of two subsidiary and two guard cells. Nuclei/DNA are shown in magenta; microtubule structures are shown in green. **(G)** Ordered sequence of protein polarization, which occurs at the time shown in panel B. (**H-K**) Mutants with polarity defects result in abnormal subsidiary cells exhibit a range of abnormal morphologies. Examples of a single stomatal complex from wild type, *brk1, pan2,* and *pan1* mutants, where abnormal subsidiaries are false-colored yellow. Supports Figures 1 and 5.

**Supplemental Figure 2.**
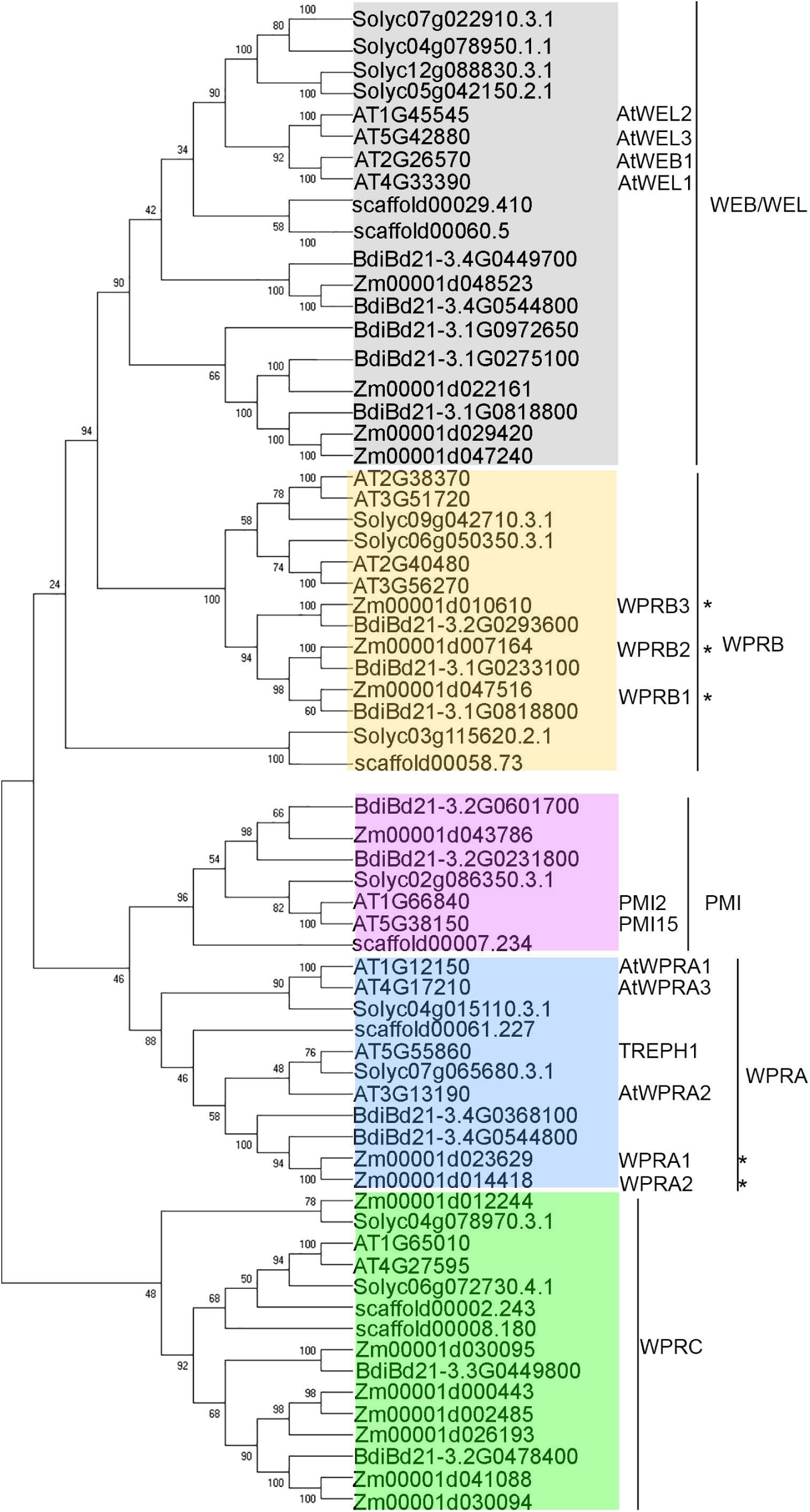
Protein tree of the WPR family in plants. Protein coding sequences of WPR family members were identified from Phytozome 13 (https://phytozome-next.jgi.doe.gov/) from *A. thaliana* (dicot), *Amborella trichopoda* (dicot), *Solanum lycopersicum* (tomato, dicot), *B. distachyon* (monocot) and *Zea mays* (maize, monocot), and a protein tree was inferred. The proteins fall into 5 subfamilies, including the previously identified WEB/WEL, WPRB, WPRA and PMI clades – we also identified an additional WPRC clade. Previously named proteins are listed, and asterisks mark proteins characterized in this study. Protein sequences were aligned using Clustal Omega and the tree was inferred using MEGA 11. Numbers at nodes represent the percentage values given by 1000 bootstrap analysis samples. At: *Arabidopsis thaliana*, Zm: *Zea mays*. Bd: *Brachypodium distachyon,* Solyc: *Solanum lycopersicum*, scaffold: *Amborella trichopoda.* Supports Figure 1.

**Supplemental Figure 3.**
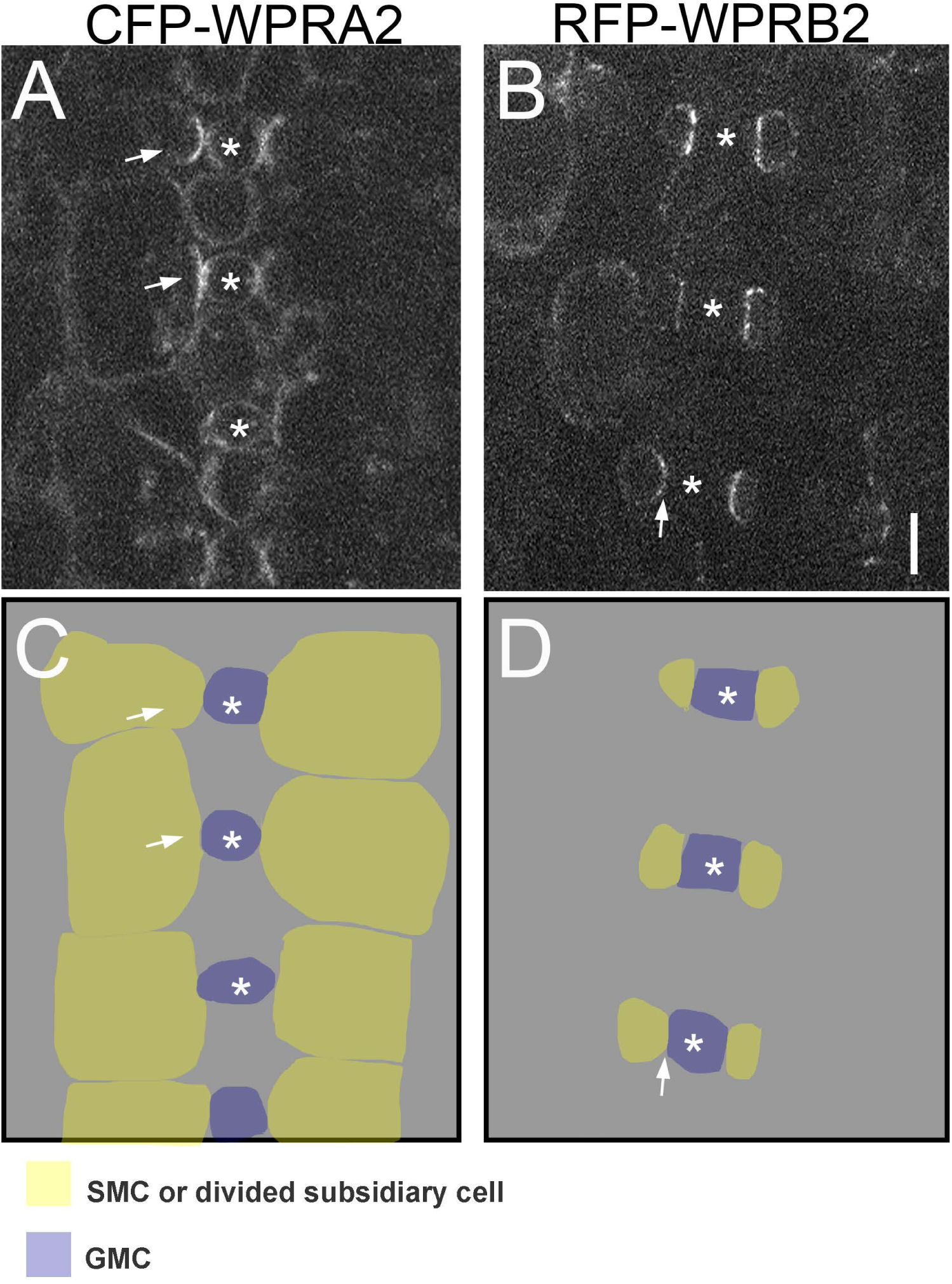
Plasmolysis assays of CFP-WPRA2 and RFP-WPRB2 transgenic plants. Plants expressing either CFP-WPRA2 or RFP-WPRB2 were treated with 0.8M mannitol to induce plasmolysis, to determine if WPR proteins were primarily localized to SMCS or GMCs. Similar results were found with both fusions, with representative images shown. Although plasmolysis results in significant loss of overall signal, fluorescence was still visible in some cells, especially in SMCs. (A) CFP-WPRA2 expressing cells. Arrows point to bright fluorescent patches in two cells that are SMCS. Based on the curvature of the membrane, away from GMCs (marked by asterisks), the polarized fluorescence is predominantly in SMCs. Residual signal could also be seen in GMCs, although typically was much stronger in SMCs. (B) RFP-WRPB2 expressing cells. In these more mature cells, the SMCs have likely already divided, and polarized fluorescence remains in subsidiary cells. Arrow points to a cell where fluorescence is clearly seen in the subsidiary cell, but not the adjacent GMC. (C) and (D) are cartoon schematics of (A) and (B) respectively, with color-coding of cell types. Scale bar = 10 µm. Supports Figure 1.

**Supplemental Figure 4.**
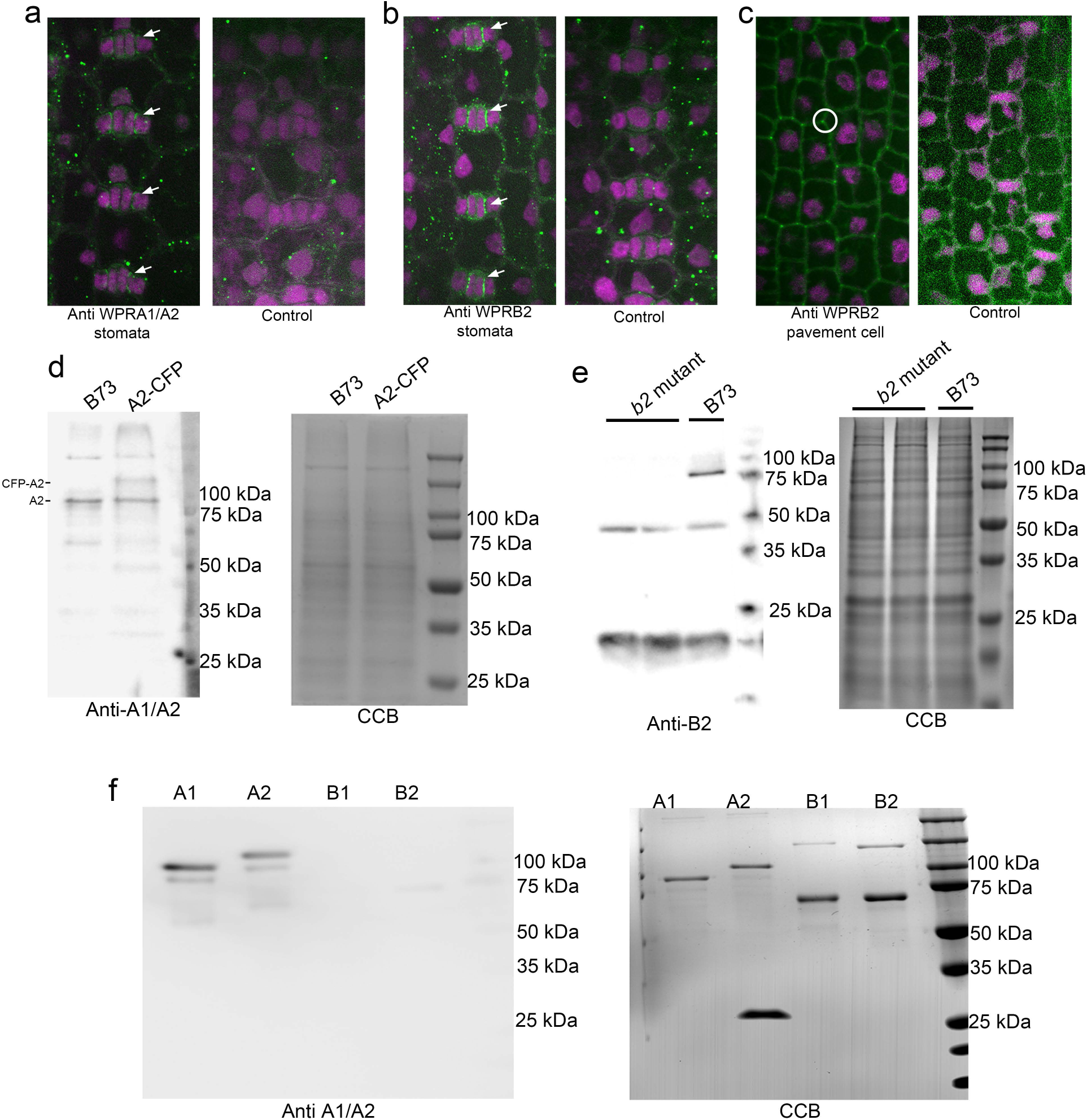
Immunolocalization using WPRA1/2 and WPRB2 antibodies and their validation. **(a)** Immunostaining with anti-WPRA1/A2 (green) antibodies and propidium iodide-staining (magenta) of developing leaf 4. No anti-WPRA1/A2 was used as a negative control. Arrows indicate the localization of WPRA1/A2 at the interface of subsidiary cells and guard cells. **(b and c)** Immunostaining with anti-WPRB2 (green) antibodies and propidium iodide (magenta). CRISPR-Cas9 induced *wprb2* mutants were used as a negative control. Arrows in (b) indicate the localization of WPRB2 at the interface of subsidiary cells and guard cells, in (c) indicate the localization of WPRB2 at pavement cell corner. **(d-f)** Confirmation of anti-WPRA1/A2 **(d and f)** and anti-WPRB2 **(e)** specificity using Western blotting. **(d)** Western blot of proteins extracted from B73 and CFP-WPRA2 transgenic plants probed with affinity-purified anti-WPRA1/A2. A band corresponding to the predicted size of endogenous WPRA1/A2 was recognized in B73. The same band was recognized in CFP-WPRA2 expressing plants; as was a larger band specific to the transgenic plants, which corresponds to the predicted size of CFP-WPRA2. In the right panel, Coomassie blue (CCB) staining of the gel is shown as a loading control. **(e)** Proteins extracted from *wprb2* mutants and B73 were used for western blotting assays. A band around 75 kD which consistent with the mass of WPRB2 protein was recognized in B73 but not in two *wprb2* mutants, Coomassie blue (CCB) staining is shown as a loading control. **(f)** Western blot of bacterially-expressed and purified GST-tagged WPRA1, WPRA2, WPRB1, WPRB2 proteins probed with affinity-purified WPRA1/A2 antibody demonstrates the specificity of this antibody for WPRA proteins. The Coomassie blue–stained gel (CCB) in the lower panel shows the purified proteins of WPRA1, WPRA2, WPRB1, WPRB2. Supports Figure 1, Figure 2 and Supplemental Dataset 2.

**Supplemental Figure 5.**
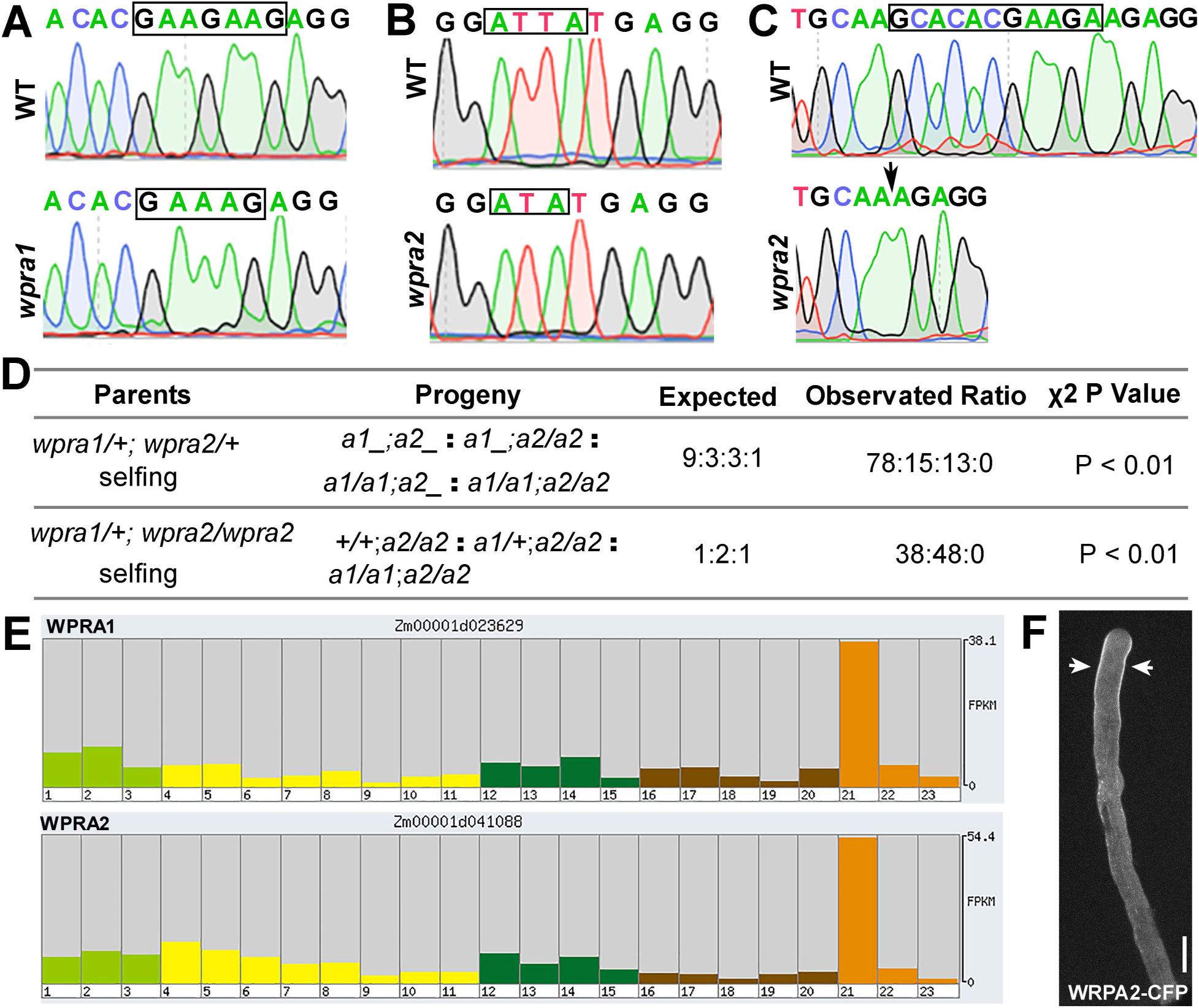
CRISPR-Cas9 induced *wpra1 and wpra2* mutants have transmission defects. Mutated regions are labeled with black squares. **(A)** DNA sequences of Cas9-edited *Wpra1*. *wpra1* has a two nucleotide “GA” deletion at the gRNA2-targeted site. **(B)** and **(C)** DNA sequences of Cas9-edited *Wpra2,* Two mutations were identified: a “T” deletion was found at the gRNA1-targeted site (B), and an 11bp (GCACACGAAGA) deletion was found at gRNA2-targeted site (arrow, C). **(D)** Genotype analysis was performed in the progeny of *wpra1*/+; *wpra2*/+ and *wpra1*/+; *wpra2*/*wpra2* mutants. No *wpra1*/*wpra1*; *wpra2*/*wpra2* double mutants were recovered; observed ratios are significantly different from expected ratios. **(E)** RNA-seq expression level in maize growth stages (Data from MaizeGDB, Walley et al., 2016). Stage 21 is B73_Mature Pollen (expression level = 54.4). **(F)** CFP-WPRA2 localization in maize pollen tube. Maize pollen stably expressing CFP-WPRA2 were germinated in liquid culture for 1 hour at room temperature observed using spinning disk confocal and images were collected at × 60 objective (NA = 1.20, Olympus). The arrows point to subapex region localization of CFP-WPRA2. Scale bar in (F) = 10 µm. Supports Figure 5.

**Supplemental Figure 6.**
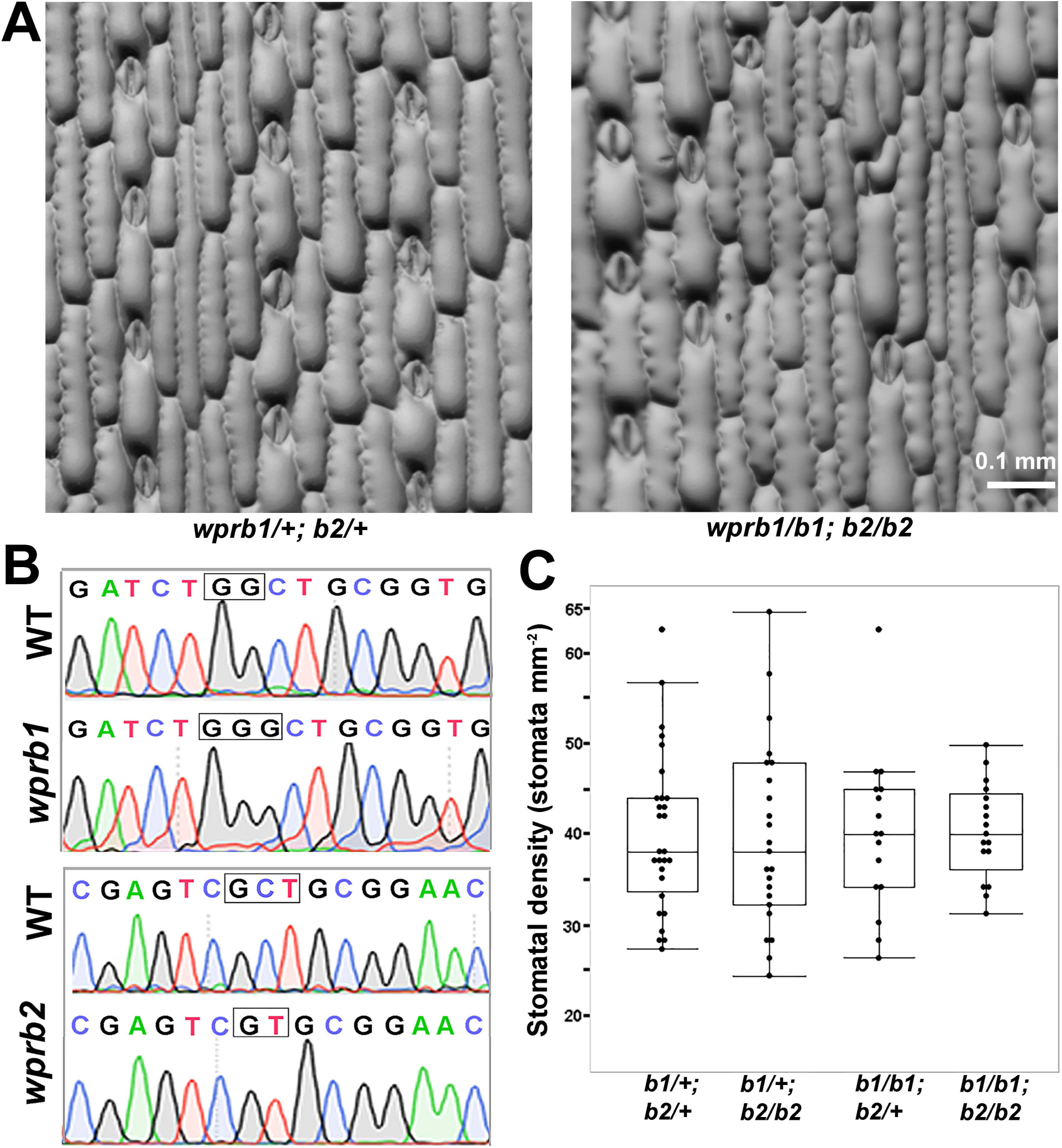
CRISPR-Cas9 induced *wprb1;wprb2* double mutants have no effect on stomatal density. **(A)** Representative image of the third leaf of epidermis of *wprb1/+;wprb2/+* and *wprb1/b1;wprb2/b2* mutants. Scale bar, 0.1 mm. **(B)** Cas9-edited *Wprb1* and *Wprb2* genes. Mutated regions are labeled in black square; wprb1 has a “G” insertion and wprb2 has a “C” deletion in the gRNA targeted region. These mutations cause frameshifts and premature termination. **(C)** Quantification of stomatal density in *wprb1/+; wprb2/+* (n = 27 plants), *wprb1/b1;wprb2/+* (n = 14 plants), *wprb1/+;wprb2/wprb2* (n = 22 plants) and *wprb1/wprb1;wprb2/wprb2* (n=18 plants). Student’s T-tests were performed among these genotypes, no significant difference in these mutants. Supports Figure 5.

**Supplemental Figure 7.**
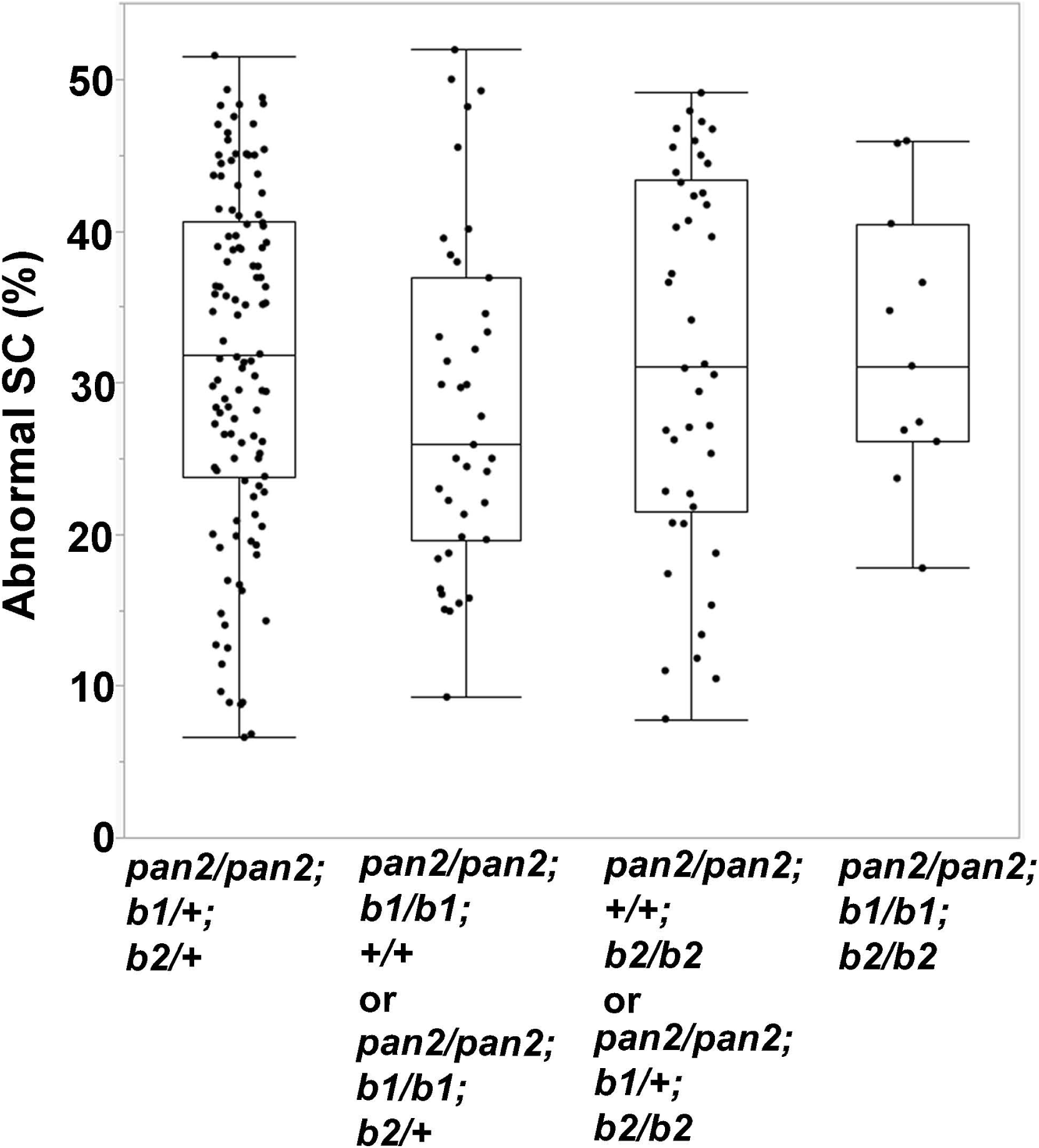
Mutations in *wprb1* and *wprb2* do not enhance *pan2*/*pan2* phenotype. Stomatal phenotypes of progeny of *pan2*/*pan2; wprb1*/*+; wprb2*/*+* plants were performed using the fully expanded third leaf. The percentage of abnormal subsidiary cell (SC) was quantified in different genotypes, *pan2/pan2; wprb1*/*+; wprb2/+* (n = 113 plants)*, pan2/pan2; wprb1*/*wprb1;wprb2_*(n = 38 plants)*, pan2*/*pan2;wprb1_; wprb2*/*wprb2* (n= 41 plants) *and triple mutants pan2*/*pan2;wprb1*/*wprb1; wprb2*/*wprb2*(n= 11 plants). For each plant 100-200 subsidiary cells were genotyped. ANOVA and pair-wise student’s T-tests were performed among these genotypes with no significant differences. Supports Figure 5.

**Supplemental Figure 8.**
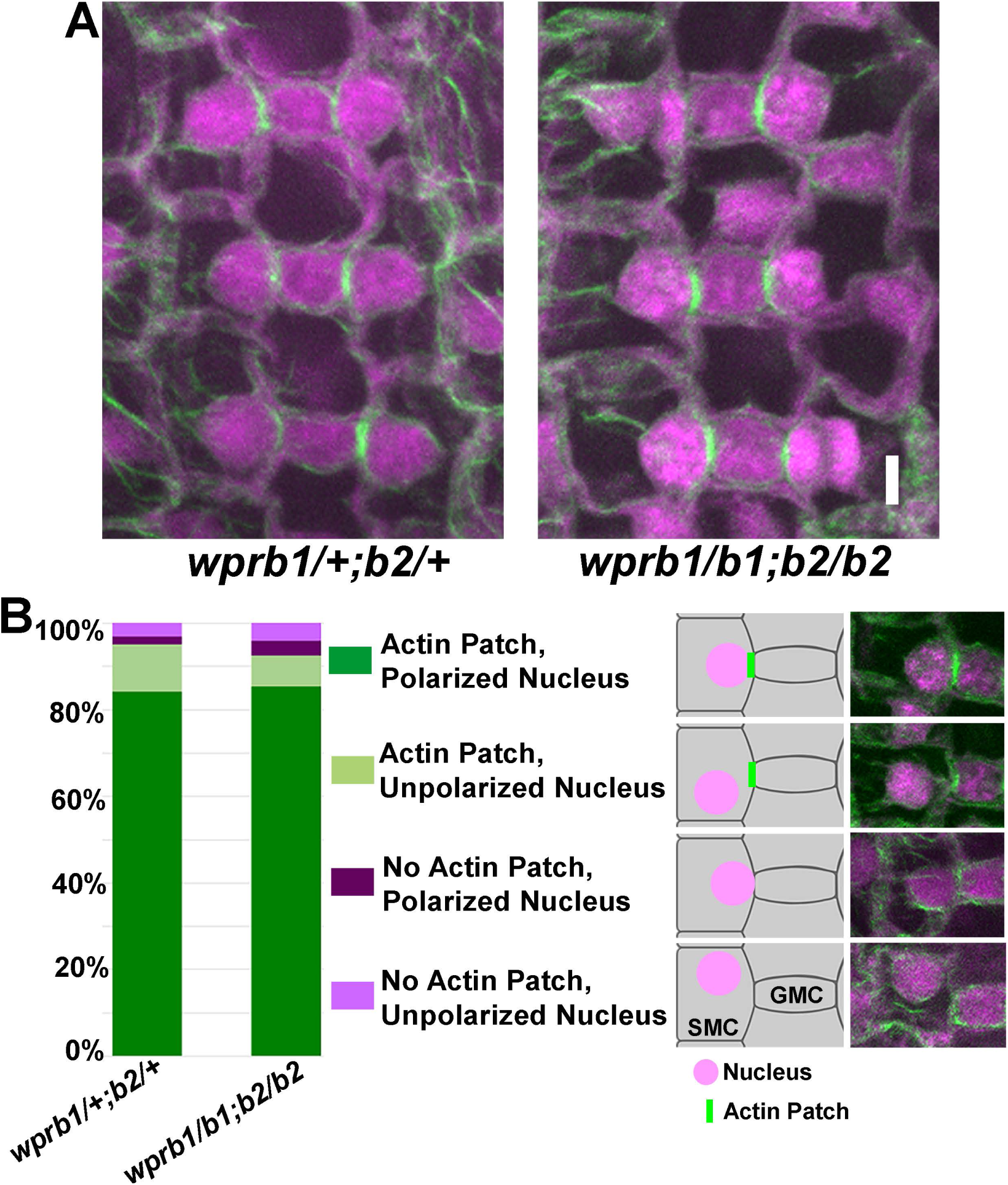
CRISPR-Cas9 induced *wprb1;wprb2* double mutants have no effect on polarized actin accumulation or nuclear polarization during maize SMC development. The stomatal division zone of developing leaf 4 from *wprb1;wprb2* double mutants, and heterozygous siblings was stained using Alexafluor488-phalloidin (green) and propidium iodide (magenta). SMCs adjacent to GMCs that were greater than than 6 µm in width were used for the quantification, at this stage the majority of SMCs in wild-type had both actin patch and polarized nucleus at SMC-GMC interface. Cells were assayed for the presence of an actin patch, and whether the nucleus was polarized (touching the SMC-GMC interface). (A) Representative image of Alexafluor488-phalloidin stained actin (green) and propidium iodided stained DNA (magenta) in leaf epidermis from *wprb1/+;wprb2/+* and *wprb1/b1;wprb2/ b2* plants. Bar = 5 µm. (B) Quantification of SMC classifications in *wprb1/+;wprb2/+* (n = 3 plants, 221 cells) and *wprb1/b1;wprb2/b2* (n = 3 plants, 241 cells) sibling plants. Supports Figure 5.

**Supplemental Figure 9.**
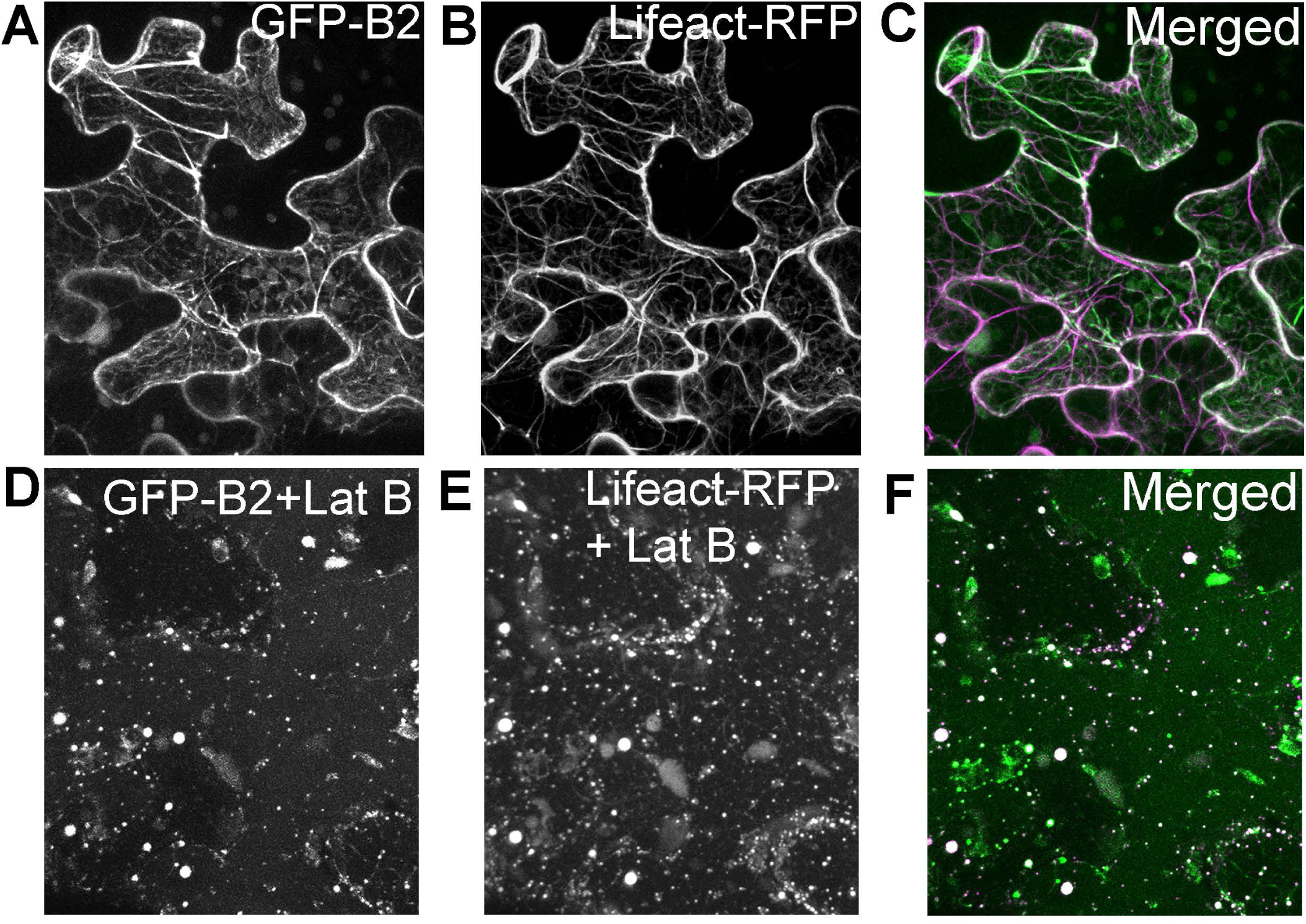
Latrunculin B-treated tobacco leaf cells transiently expressing GFP-WPRB2 and Lifeact-RFP. Tobacco leaves co-expressing GFP-WPRB2 and Lifeact-RFP were treated with DMSO (negative control, **A-C)** or 40 μM Latrunculin B **(D-F)** for 2 hours. Supports Figure 6.

**Supplement Figure 10.**
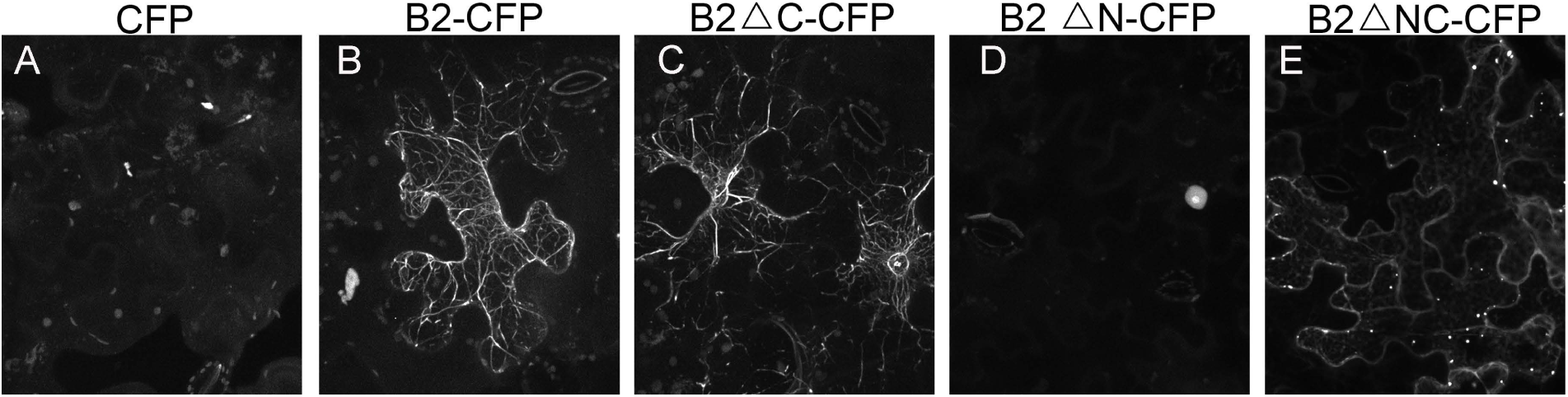
Transient expression of WPRB2-CFP, and its truncated proteins. Localization of WPRB2 in tobacco was confirmed using a CFP (rather than GFP) tag, and a C-terminal (rather than N-terminal) fusion. See Figure 6A for cartoon of protein truncations tested. (A) Soluble CFP. (B) Full length WPRB2-CFP. (C) C-terminal truncation WPRB2ΔC-CFP (D) N-terminal truncation WPRB2ΔN-CFP localizes predominantly to the nucleus. (E) Central DUF827 domain only WPRB2ΔNC. Filaments structures appears in WPRB2-CFP and WPRB2△C-CFP expressing tobacco cells, but not in WPRB2ΔN-CFP and WPRB2ΔNC-CFP expressing tobacco cells. Supports Figure 6.

**Supplemental Figure 11.**
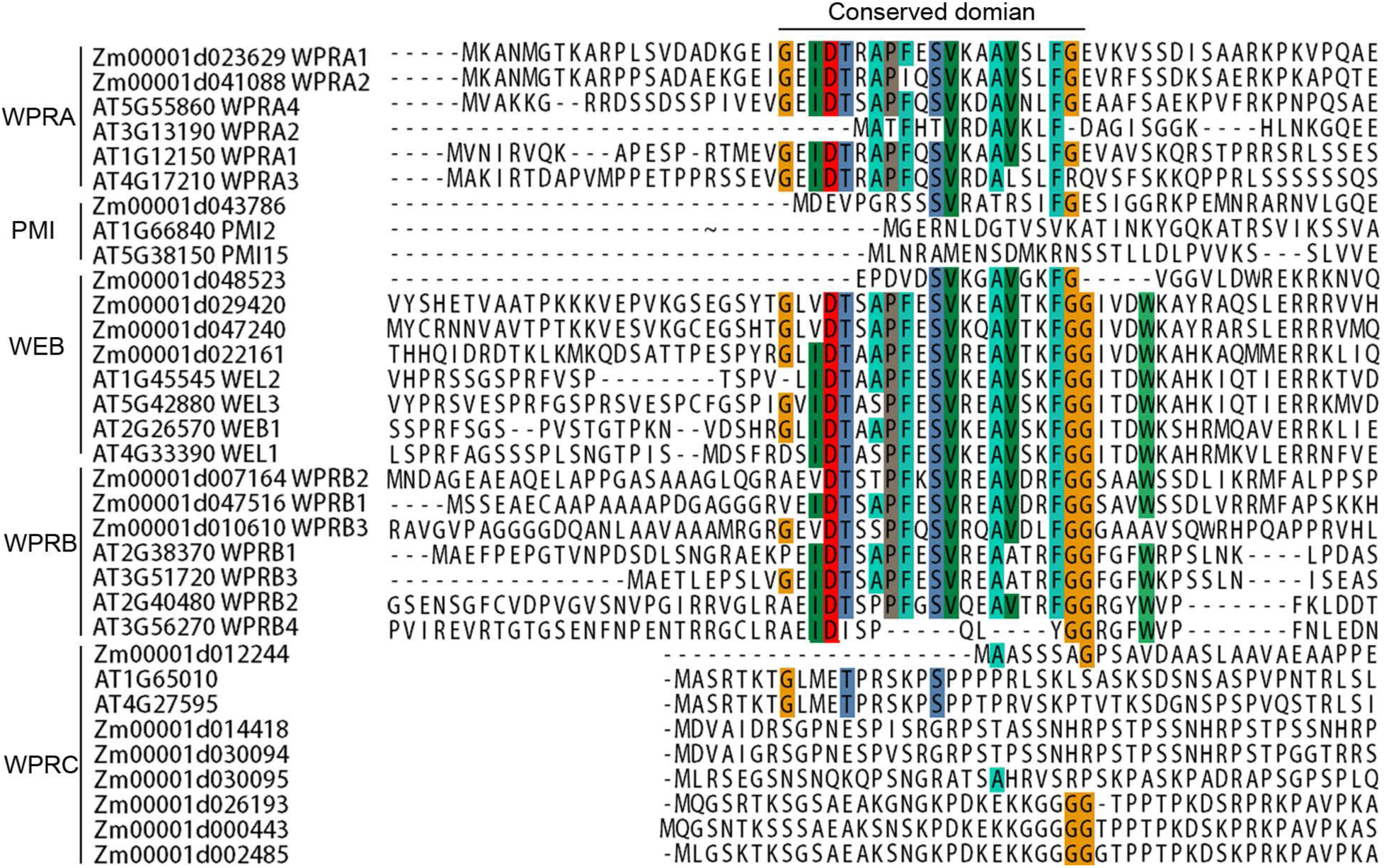
Alignment of the actin-binding domain of the WPR protein family. The N terminal amino acids sequence WPRs from *Arabidopsis* and maize were aligned using ClustalX 2.0.5 (Larkin et al., 2007) with the default settings. Supports Figure 6.

**Supplemental Figure 12.**
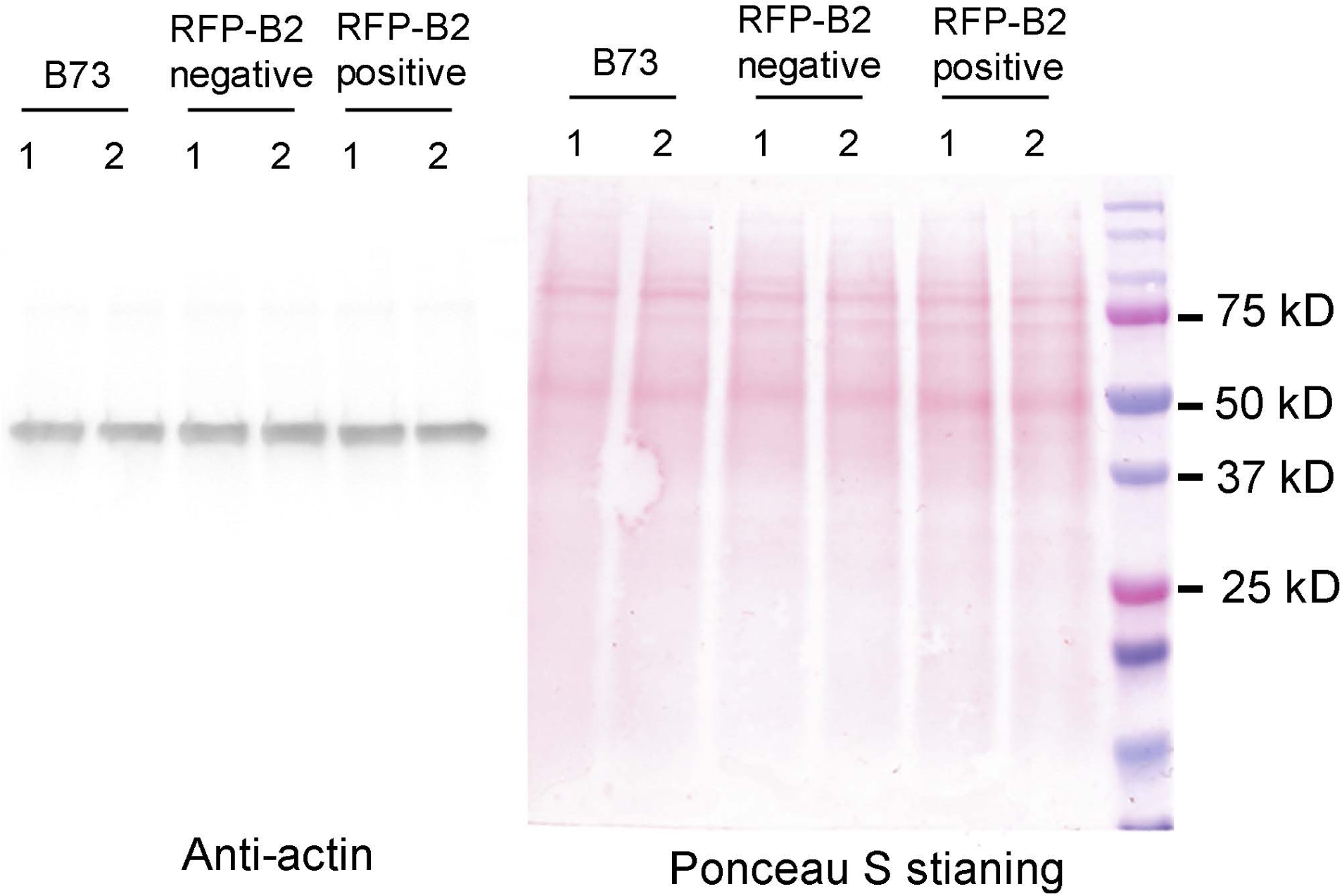
RFP-WPRB2 expression has no effect on the total amount of actin. Western blot of total proteins extracted from the stomatal division zone of two independent plants of B73, RFP-WPRB2 transgenic plants, RFP negative siblings. Extracts were probed with an anti-actin antibody. Ponceau S staining is shown as a loading control.

**Supplemental Figure 13.**
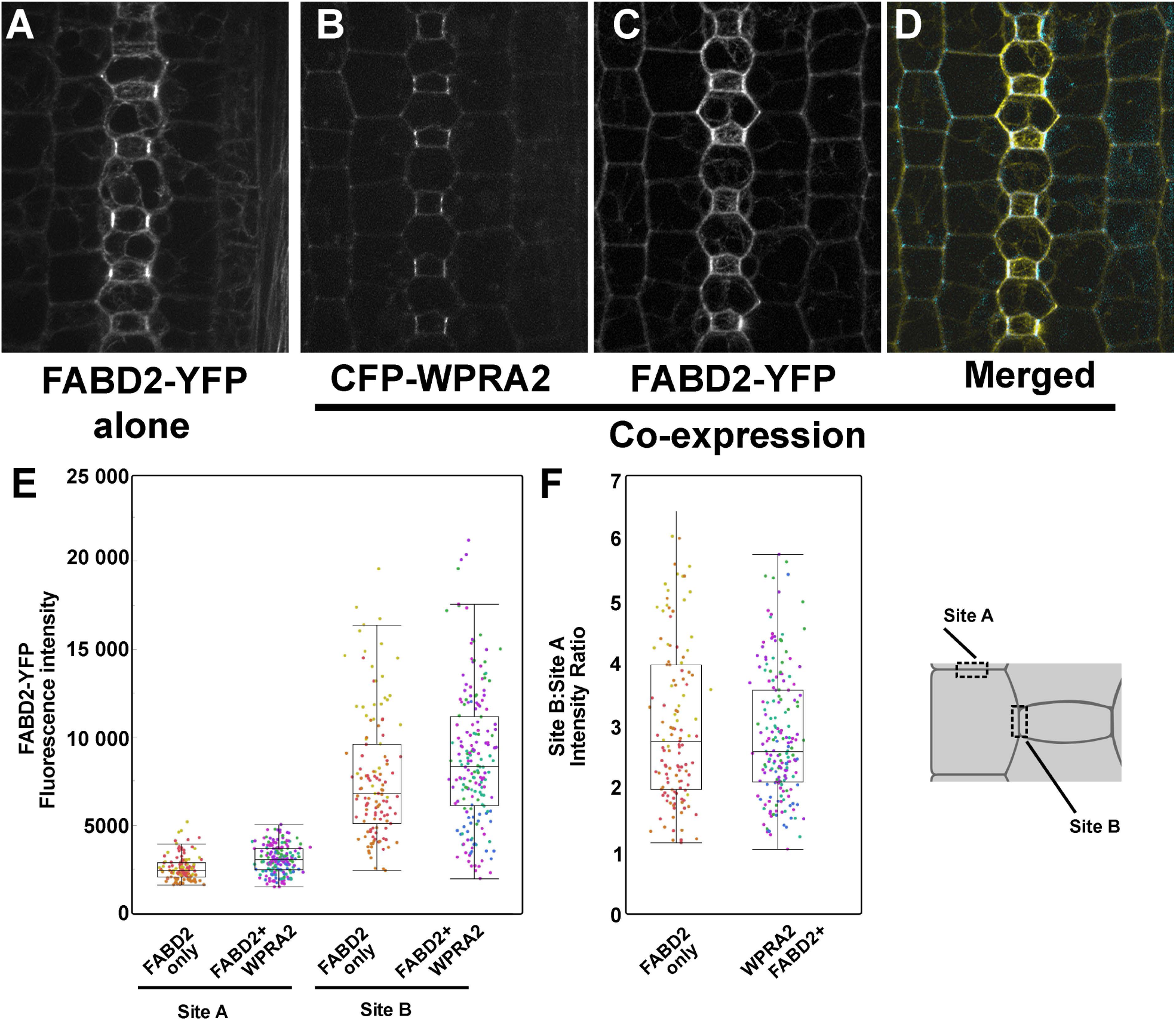
Co-expression of CFP-WPRA2 with ABD2-YFP does not decrease FABD2-YFP intensity. Plants expressing CFP-WPRA2 and FABD2-YFP were crossed, and the progeny independently segregated the two markers. **(A)** Representative image of plant only expressing FABD2-YFP. **(B-D)** Plant co-expressing FABD2-YFP (yellow in D) **(C)** with CFP-WPRA2 (cyan in E) and the merged image **(D),** Scale bar in (D) = 5 µm. **(E, F)** Quantification of fluorescence intensity and intensity ratio of SMC lateral cell side (site A) and SMC-GMC interface (site B) in FABD2-YFP only plants (n = 3 plants, 254 cells) and CFP-WPRA2 + FABD2-YFP co-expressing plants (n = 5 plants, 379 cells). Images are 7-slice max projections. Dots of the same color are from the same plant. Supports Figure 7.

**Supplemental Figure 14.**
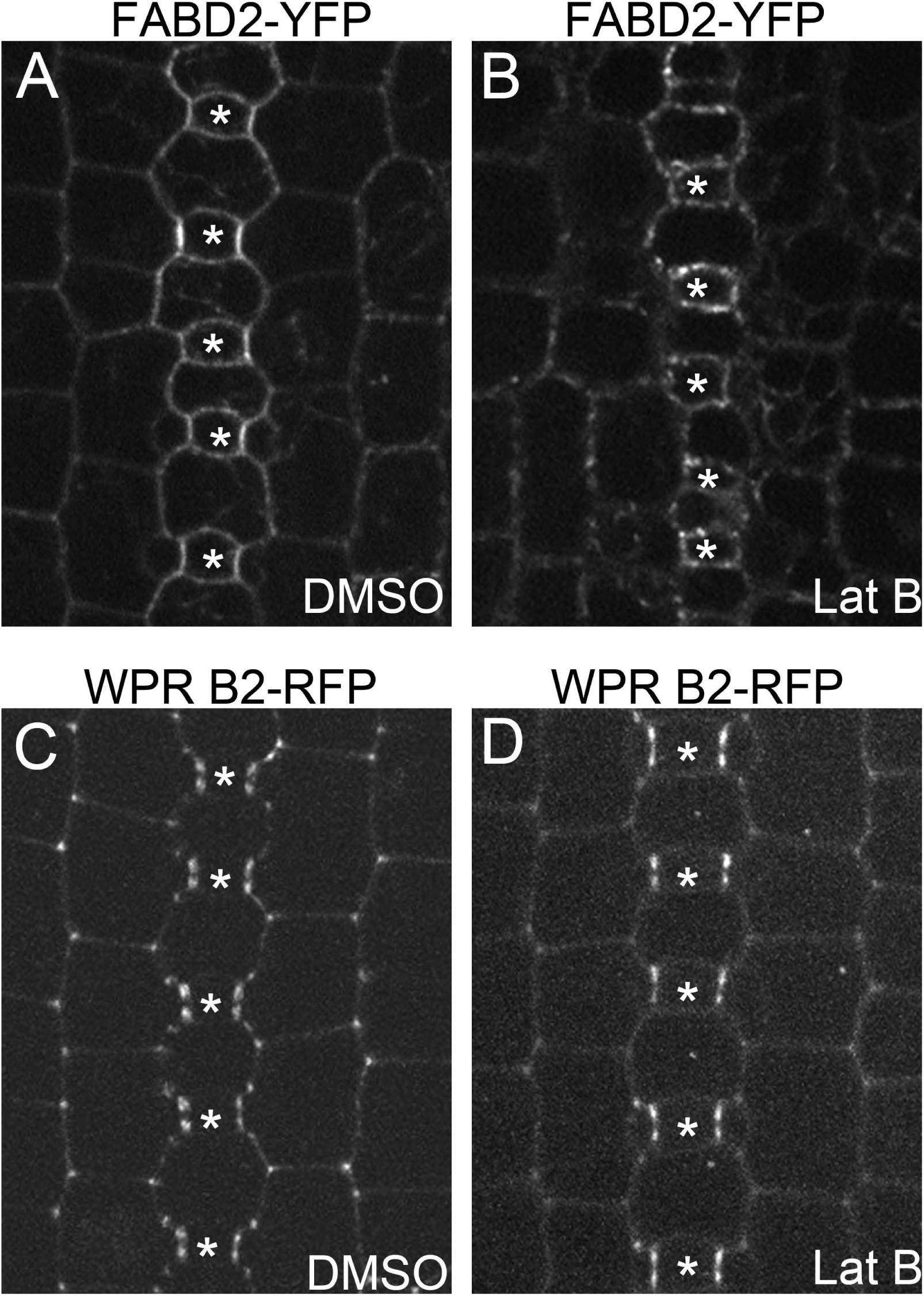
F-actin disruption does not affect the polarized localization of RFP-WPRB2 in SMCs. **(A)** FABD2-YFP expressing maize leaves were treated with DMSO (negative control) and 40 μM Latrunculin B for 4 hours. **(B)** RFP-WPRB2 expressed maize leaves were treated with DMSO (negative control) and 40 µM Latrunculin B for 4 hours. Asterisks mark GMCs.

**Supplemental Data Set 1. Co-IP/MS results using an anti-GFP antibody and membrane extracts from PAN2-YFP, PAN1-YFP, BRK1-CFP, PDI-YFP, PIN1-YFP, Rab11D-YFP and B73**

**Supplemental Data Set 2. Co-IP/MS results using WPRA1/2, WPRA2-CFP, WPRB2-RFP as bait (and corresponding controls)**

**Supplemental File1. Amino acid sequences of WPRs used for the phylogenetic analysis in Supplemental Figure 2.**

## Notes

### Competing Interest Statement

The authors have declared no competing interest.

### Summary of Updates

Addition of co-IP/MS data for PAN2-YFP, and addition of two authors that conributed to those experiments. Relabelling/reordering/updating panels of figures 3, 5, 6, and 7. Addition of 4 new supplemental figures. No changes in the major conclusions, however supporting data was added.

